# Vaccine Elicitation of HIV-1 Neutralizing Antibodies Against Both V2 Apex and Fusion Peptide in Rhesus Macaques

**DOI:** 10.1101/2025.09.29.679292

**Authors:** Hongying Duan, Joseph P. Nkolola, Shuishu Wang, Jayeshbhai Chaudhari, I-Ting Teng, Christy Lavine, Danealle K. Parchment, George S. Sellers, Krisha McKee, Sijy O’Dell, Misook Choe, Haijuan Du, Baoshan Zhang, Alejandro A. Espinosa Perez, Annika Rossler, Ninaad Lasrado, Andrea Biju, Jordan E. Becker, Robin Carroll, Audrey S. Carson, Amy R. Henry, Nicholas C. Morano, Madeeha Mughal, Reda Rawi, Ryan S. Roark, Chaim A. Schramm, Chen-Hsiang Shen, Sarah C. Smith, Tyler Stephens, Yaroslav Tsybovsky, David J. Van Wazer, Hua Wang, Yongping Yang, Lucy Rutten, Johannes P.M. Langedijk, Cheng Cheng, Lingshu Wang, Daniel C. Douek, Richard A. Koup, John R. Mascola, Lawrence Shapiro, Tongqing Zhou, Nicole A. Doria-Rose, Bette Korber, Michael S. Seaman, Theodore C. Pierson, Peter D. Kwong, Dan H. Barouch

## Abstract

Broadly neutralizing antibodies targeting multiple sites of HIV-1 Env vulnerability can be induced by infection, but simultaneous elicitation of neutralizing antibodies (NAbs) against multiple epitopes has not yet been achieved by vaccination. In this study, we designed a dual-epitope vaccine targeting both fusion peptide (FP) and V2 apex and evaluated its capacity to induce NAbs against both epitopes in rhesus macaques. This vaccine combined an FP conjugate with a cocktail of engineered Env trimers with enhanced V2 apex recognition and increased antigen retention in lymph nodes. Immunization of macaques with the dual-epitope vaccine elicited >1000-fold higher autologous tier 2-neutralization titers than the wildtype Env trimer and enhanced heterologous NAb breadth. Both FP and V2-apex monoclonal antibodies (mAb) were isolated from immunized macaques and showed heterologous neutralization with genetic and structural signatures that were similar to well-characterized FP and V2 apex bNAbs, although the V2 apex mAbs showed incomplete maturation. These results demonstrate proof-of-concept for simultaneous vaccine elicitation of NAbs against multiple sites of Env vulnerability, which will likely be critical for an effective HIV-1 vaccine.

**HIGHLIGHTS:**

- Designed a dual-epitope vaccine targeting both fusion peptide (FP) and V2 apex
- V2-SET Env trimer conferred higher binding of V2 apex bNAbs, longer retention in draining lymph nodes, and >1000-fold higher induction of autologous neutralization titers compared with wildtype Env trimer
- Dual-epitope vaccine enhanced serum tier 2 neutralization breadth
- Dual-epitope vaccine elicited FP– and V2 apex-specific neutralizing mAbs with genetic signatures similar to well-characterized FP and V2 apex bNAbs

## INTRODUCTION

Although HIV-1 was identified over four decades ago, a protective vaccine against HIV-1 remains elusive. The HIV-1 envelope (Env) glycoprotein trimer is the only target of virus-specific neutralizing antibodies (NAbs), making it an attractive focus for vaccines. Progress towards an effective HIV-1 vaccine has been hindered by the high rate of mutation and recombination during viral replication (Bbosa et al., 2019; Ferguson et al., 2002), a dynamic Env structural ensemble (Cale et al., 2022; Hodge et al., 2022; Munro and Lee, 2018), and the extensive glycan shield of ∼90 N-linked oligosaccharides (Stewart-Jones et al., 2016). Soluble Env trimers from multiple HIV-1 strains have been stabilized in prefusion-closed conformations (Sanders et al., 2013), but immunization with stabilized soluble trimers has generally failed to elicit robust heterologous neutralizing titers (Bianchi et al., 2018; Corrigan et al., 2021; Cottrell et al., 2020; Hu et al., 2015).

Several NAb epitopes on HIV-1 Env have been targets for vaccine development, including the fusion peptide (FP), V2 apex, V3-glycan supersite, and CD4-binding site. Current efforts in the development of an HIV-1 vaccine have typically focused on induction of NAbs to a single target epitope (Martinez-Murillo et al., 2017; Mogus et al., 2020; Pratap et al., 2025; Voss et al., 2017; Wang et al., 2024; Xu et al., 2018), but an effective HIV-1 vaccine will almost certainly need to target multiple sites of Env vulnerability given the challenge of HIV-1 diversity. Previously, bNAbs targeting multiple epitopes have been described in persons with HIV-1 (Bonsignori et al., 2012; Cale et al., 2017; Cale et al., 2024; Cheng et al., 2019; Huang et al., 2014; Huang et al., 2012; Setliff et al., 2019), but multi-epitope targeting vaccines have not yet been developed (Gristick et al., 2025).

We previously reported that priming with FP-carrier conjugates followed by boosting with soluble prefusion-closed stabilized Env trimers elicited FP-specific bNAbs in nonhuman primates with up to 59% neutralizing breadth as assessed on a cross-clade panel of 208 HIV-1 strains (Kong et al., 2019; Kong et al., 2016; Xu et al., 2018). In addition, recombinant tetanus toxoid heavy chain fragment (rTTHC) linked to FP8 (FP8-rTTHC) was evaluated in animal models as a suitable FP-conjugate vaccine immunogen (Ou et al., 2020) and has recently been advanced into a phase I clinical trial. We also previously defined a V2 bNAb signature associated with bNAb sensitivity and resistance, and we selected and modified the V2 sequence of acute clade C isolate 459C to create V2 signature-based epitope-targeted (V2-SET) 459C trimers, with the 459C Optimal (OPT) trimer modified with V2 bNAb sensitive mutations and 459C Alternative (ALT) trimer modified to capture natural diversity within V2, which included signatures of resistance. A trimeric cocktail of 459C WT, OPT, and ALT trimers comprised the 459C V2-SET Env vaccine, and immunization with this trimeric cocktail conferred enhanced neutralizing breadth and potency as compared to the 459C wildtype (WT) Env vaccine in guinea pigs (Bricault et al., 2019).

In this study, we explored the potential of combining FP8-rTTHC immunogens and 459C V2-SET Env trimers to induce NAbs to both FP and V2 apex sites of vulnerability. We optimized the V2-SET Env trimer stabilization with a repair and stabilization approach for a study in rhesus macaques, and we evaluated sera and isolated mAbs that targeted both epitopes. Our data demonstrate simultaneous induction of NAbs against both sites of vulnerability with genetic and structural signatures of well-characterized bNAbs.

## RESULTS

### V2-OPT modifications increase binding to V2 bNAbs and capture and retention in lymph nodes and spleens of mice

Previously reported viral signatures associated with V2 apex-specific bNAb sensitivity and diversity were employed to create the 459C V2-SET vaccine (Bricault et al., 2019; Rutten et al., 2018), which is a set of three trimers designed such that, in combination, the most common amino acid variants within the V2 epitope region in the globally circulating viral population are covered. The cocktail included a 459C wildtype (WT) trimer, a natural strain that showed intrinsic capacity to elicit NAbs with some heterologous breadth; a 459C optimized (OPT) trimer, which included amino acid modifications that represented common variants not found in 459C and that were also associated with containing a V2 apex-specific bNAb sensitivity sequence signatures; and a 459C alternate (ALT) trimer, which included modifications that provided further coverage of the natural diversity of HIV-1 sequences within the V2 epitope (Figures 1A, S1A). HIV-1 Env hypervariable V1 and V2 region characteristics known to be important for V2 apex bNAb sensitivity (short loops that are positively charged) were also factored into the 459 V2-SET design (Bricault et al., 2019). Stabilizing the prefusion conformation is important to reduce the exposure of non-neutralizing epitopes and to elicit a more focused immune response towards neutralizing epitopes (de Taeye et al., 2015; Kulp et al., 2017) and therefore we incorporated Repair-and-Stabilize (RnS) mutations (Rutten et al., 2018) into the sequences of the 459C WT, OPT, and ALT trimers (Figures 1A, S1B).

**Figure 1.**
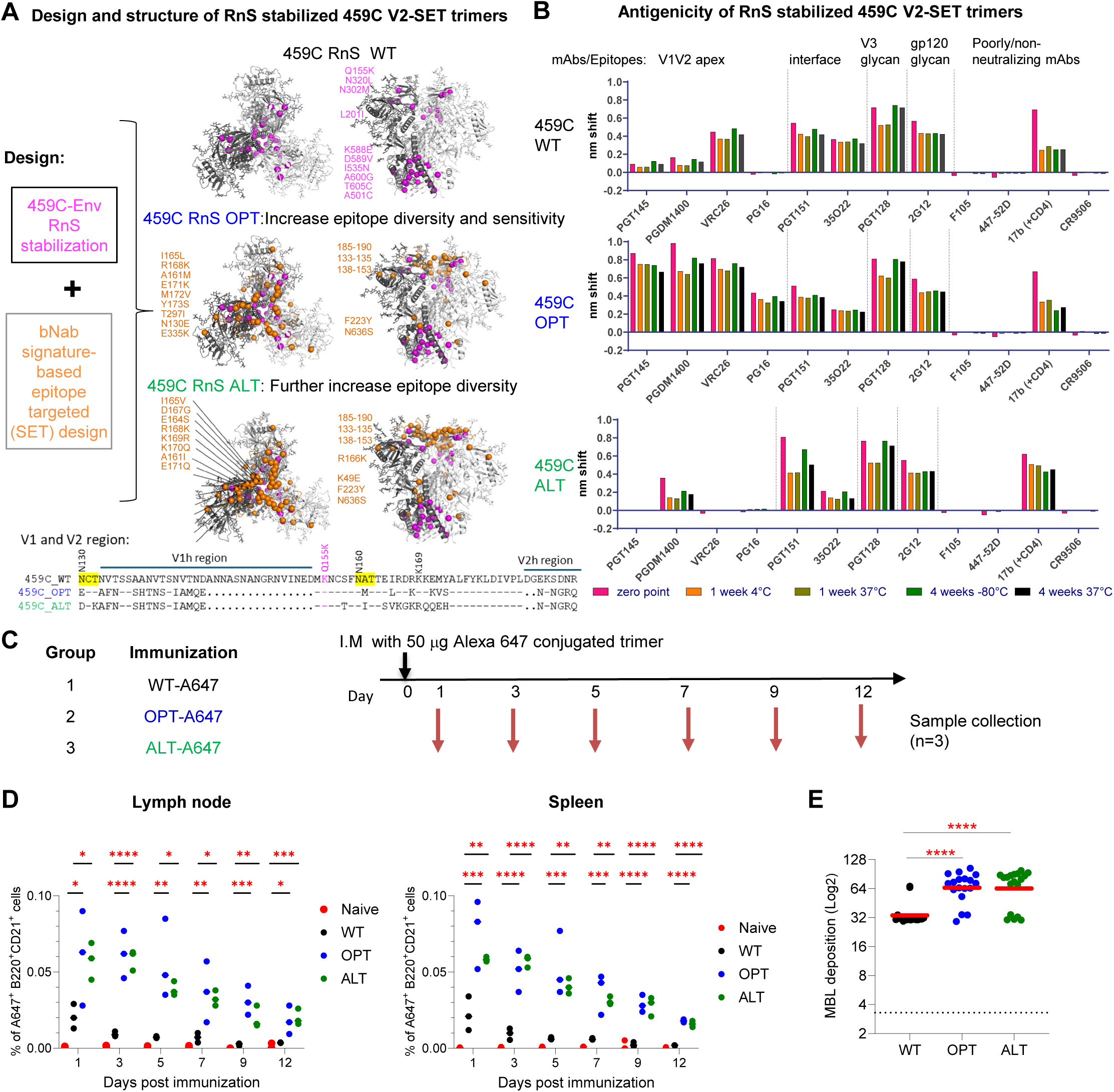
Design and structure of RnS V2-SET Env trimers that bind V2-apex bNAbs and are more efficiently captured and retained in lymph nodes and spleens of immunized mice. (A) Designs combining RnS stabilizations and V2 SET modifications for 459C WT, OPT and ALT trimers and their structures determined by cryo-EM (Data S1). RnS and V2-SET mutations are highlighted with purple and orange spheres, respectively; RnS stabilization mutations on all three trimers are labeled on the cryo-EM structure of WT-RnS (top panel, left: top view, right: side view). OPT mutations are labeled on the WT-RnS structure (middle panel). ALT mutations are labeled on the ALT-RnS trimer structure (bottom panel). Amino acid alignment in V1V2 region are shown below the cryo-EM structure panels, with glycans at 130 and 160 highlighted in yellow, RnS mutant Q155K shown in purple, and V2-bNAb core recognition site K169 shown on the top of the alignment. V1h: V1 hypervariable region; V2h: V2 hypervariable region. (.): missing residue. (−): identical residue. (B) Binding of 459C WT, OPT and ALT trimers to HIV-1 specific mAbs targeting different epitopes, detected by BLI. (C) Immunization regimen of RnS 459c WT, OPT and ALT trimers in Balb/C mice. 459C RnS WT, OPT and ALT trimers were conjugated with Alexa 647 and injected I.M. in Balb/C mice. Three animals from each group were euthanized on indicated days post-immunization to harvest the spleen and lymph nodes. (D) Frequency of A647+cells in total B220+CD21+ cells in draining lymph nodes (left) or spleen (right) were evaluated with FACS analysis. Cells were stained with anti-CD3 (T cell), anti-B220 (B cell), anti-CD21/CD35 (FDC, and B cells), and Anti-CD169 (macrophages) markers. (E) V2-SET WT, OPT, and ALT trimers were evaluated for mannose binding lectin (MBL) binding using Balb/c naive mouse serum. One-way ANOVA tests were used for statistical analysis to assess p values for different groups. *: p<0.05; **: p <0.01; ***: p <0.001; ****: p <0.0001. See also Figure S1 and Data S1.

All three trimers were expressed and purified with high yields and retained prefusion conformations at 37°C for up to 4 weeks (Figure S1C-D). Furthermore, the stability of all three trimers revealed minimal antigenic disruption after storage at –80°C, 4°C, or 37°C for 4 weeks (Figure 1B). In BLI binding assays, the 459C WT trimer showed little reactivity to V2 apex bNAbs PGT145, PGDM1400, and PG16, and moderate reactivity to CAP256/VRC26.25. As expected, the 459C OPT trimer, which included V2 antibody sensitivity signatures, showed higher binding to V2 apex-specific bNAbs PGT145, CAP256/VRC26.25, PGDM1400, and enabled binding to PG16 (Figure 1A, B). Key signature differences incorporated into 459C OPT that were associated with enhanced V2 bNAbs sensitivity included shorter more positively charged V1 and V2 loops and the elimination of the N-linked glycosylation site at position N130. In contrast, the 459C ALT trimer was less efficiently bound by V2 apex-bNAbs as anticipated by the particular V2 sequence modifications in the C strand within the V2 epitope binding region that were introduced to incorporate additional common variant amino acids into the vaccine that would need to be tolerated by vaccine-induced bNAbs to achieve breadth (Figure 1A, B). All three trimers showed similar binding to gp120/gp41 interface, FP and V3-glycan antibodies, gp120 glycan and CD4bs antibodies, and little binding to non-neutralizing F105 and 447-52D mAbs, indicating that they were folded properly with little impact on the rest of the trimer structure (Figures 1B, S1D).

To characterize the effects of the mutations on the structural features of the trimers, we determined cryo-EM structures of the individual 459C WT, OPT, and ALT RnS trimers. All three trimers were flash frozen, and single particle cryo-EM data were obtained using a Titian Krios electron microscope. We obtained a structure of the 459C WT RnS trimer at 3.3 Å and a structure of the 459C ALT RnS trimer at 3.9 Å (Data S1.1, S1.2, S1.3, S1.8). We failed, however, to obtain an interpretable map for the 459C OPT RnS trimer as the particles appeared heterogeneous. Multiple classes of 3D reconstructions with partially disordered trimers suggested the 459C OPT RnS trimer structure, while intact, was more flexible and dynamic (Data S1.1). Overall, structures of the 459C ALT RnS and 459C WT RnS trimers were similar, with a rmsd of 1.6 Å for 1586 aligned Cα atoms. Additionally, the WT and ALT structures were most similar to each other in the gp41 core helices, but shifted substantially in the V1V2 region, so that the 459C ALT RnS structure expanded slightly outwards (Data S1.1D, E). These data suggest that, in the absence of antibody binding, the 459C OPT RnS trimer is flexible whereas the 459C WT RnS trimer is more stable in a prefusion closed structure than the 459C ALT RnS and 459C OPT RnS trimers.

To investigate the impact of the V2-SET modifications in vivo, we conjugated the V2-SET trimers with an Alexa 647 fluorophore and immunized BALB/c mice with 50 μg of Alexa-647-labeled WT, OPT or ALT trimer in three groups (Figure 1C). After a single immunization, we collected spleen and draining lymph nodes from three mice per group on days 1, 3, 5, 7, 9, 12, and measured Alex-647 positive populations of B220+CD21+ cells in the lymph nodes and spleen. We found that both OPT and ALT trimers demonstrated significantly higher associations with B220+CD21+ cells in lymph nodes and spleens than WT trimers. Furthermore, OPT and ALT trimers remained detectable on B220+CD21+ cells up to day 12 after immunization, whereas the WT trimer signal declined to baseline levels by day 7. These findings suggest that V2-SET modified trimers exhibit higher capture and longer retention on follicular dendritic cells (FDC) and B cells (Figure 1D). To elucidate further the enhanced antigen capture and retention of the OPT and ALT trimers, we evaluated these three trimers with mannose binding lectin (MBL). MBL binding to glycosylated vaccine is a relevant aspect of vaccine evaluation and optimization, as binding to MBL can enhance antigen-specific germinal center (GC) responses and serum antibody titers, leading to improved vaccine efficacy (Read et al., 2022). The 459C OPT and ALT trimers showed significantly higher MBL binding than did the WT trimer (Figure 1E), which is consistent with the increased antigen capture and retention in lymph nodes and spleens.

### V2-SET Env vaccine induces >1000-fold higher autologous tier 2 neutralizing activity and greater heterologous tier 2 neutralizing activity than wiltype Env vaccine in rhesus macaques

Next, we tested the dual-epitope immunization strategy in rhesus macaques by combining FP8-rTTHC with the trivalent 459C V2-SET vaccine, comprised of 459C WT, OPT, and ALT trimers in equal amounts. Of note, 459C WT, OPT and ALT trimers all contained the same FP8 sequence as the FP8-rTTHC and exhibited similar binding to FP-specific bNAbs VRC34 and DFPH (Figure 2A). Three groups of rhesus macaques were immunized with FP8-rTTHC + 459C WT trimer (Group 1), FP8-rTTHC + 459C V2-SET trimers (Group 2), and 459C V2-SET trimers alone (Group 3), followed by six boosts with each respective trimer. At week 112, a similar immunization series was repeated (Figure 2A). We tested sera for neutralization and detected a significant difference between WT 459C and V2-SET 459C immunized animals in terms of serum neutralization titers against autologous tier 2 459C WT virus. Remarkably, at week 68, V2-SET immunized animals in G2 and G3 had median neutralizing ID50 titers of 1:149,206 and 1:213,815, respectively, with some titers >1:1,000,000. In contrast, FP + 459C WT immunized animals showed a median ID50 titer of 1:148. Similar results were observed with sera obtained at week 132.

**Figure 2.**
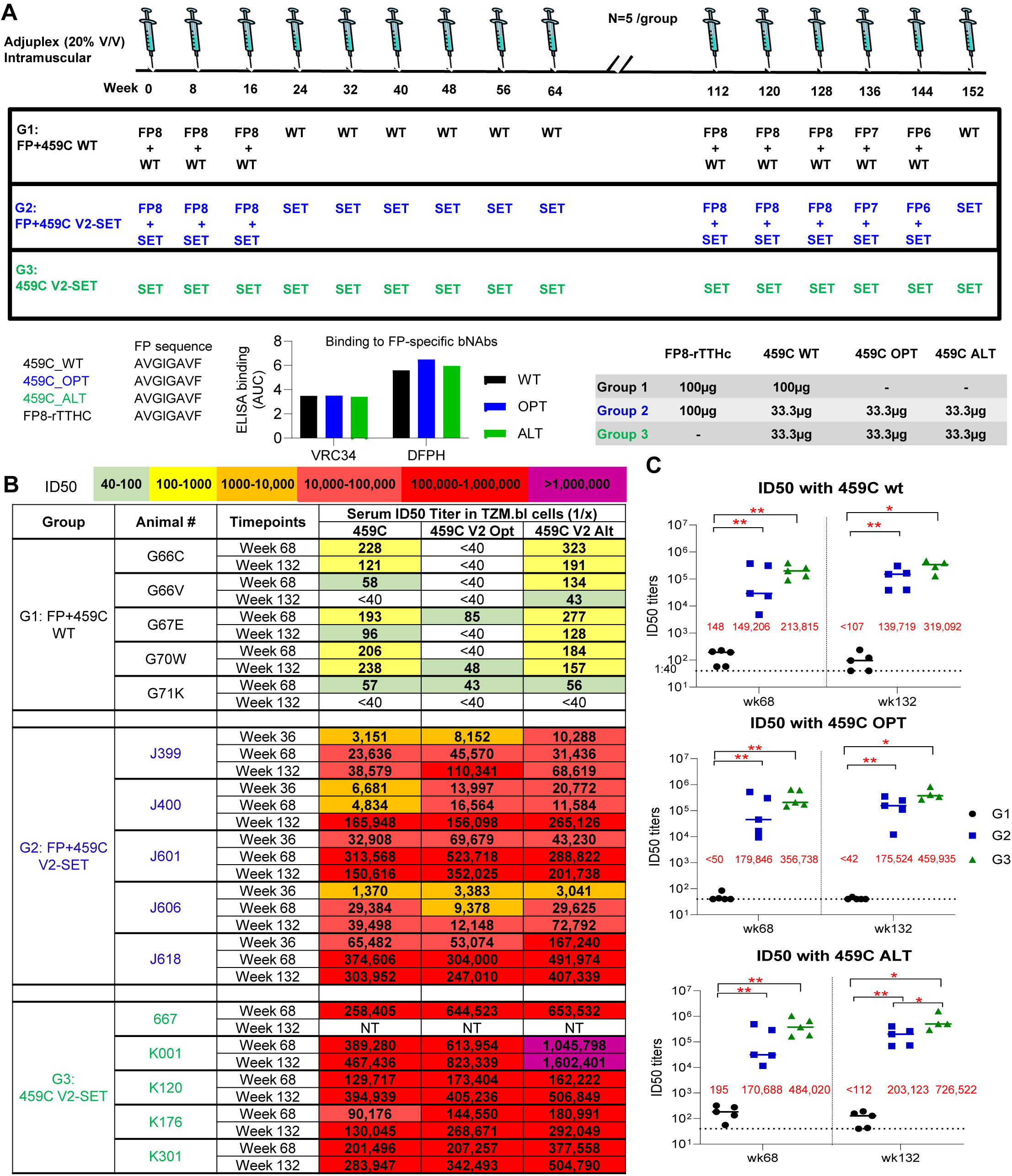
V2-SET immunization elicited 1000-fold higher autologous tier 2 neutralizing activity compared to WT trimer in all immunized NHPs. (A) Immunization schema of 3 groups of NHPs immunized with FP+459C WT trimer, FP+V2 SET trimer and V2 SET trimer alone, followed with respective trimer boost. FP-rTTHC, 459C WT, OPT and ALT contain the same FP8 sequence, and 459C WT, OPT and ALT bind to FP-specific bNabs similarly. (B,C) Plasma neutralizing ID50 titers at selected time points for vaccine-matched 459C WT, OPT and ALT viruses, which contain the V2 sequence matching the WT, OPT and ALT trimers in the immunization. (B) Color coded ID50 titers. (C) ID50 titers at wk68 and wk132 on 459C WT, OPT and ALT, respectively are shown as scatter dot plots with line at median. Median ID50 titers are shown as red fonts in the figures. Each plot represents one individual NHP in each group. Two-tailed Mann-Whitney nonparametric tests were used for statistical analysis; p-values for different comparisons are indicated *: p<0.05; **: p <0.01.

These findings demonstrate that the 459C V2-SET Env vaccine elicited autologous tier 2 neutralizing antibody titers >1000-fold greater than did 459C WT Env trimers. In addition, earlier and higher neutralizing activity against pseudotype viruses incorporating 459C OPT and ALT trimers was observed (Figure 2B-C). Interestingly, although G2 and G3 groups were immunized with only one-third of the dose of 459C WT trimer as compared to G1, sera from these animals showed far greater neutralization activity against WT 459C pseudovirus than did sera from animals immunized with WT 459C trimers only. In addition to higher autologous neutralization, 459C V2-SET immunization also elicited earlier, higher, and broader heterologous tier 2 neutralizing activity, with heterologous neutralization detected as early as week 21 (Figure S3A), including the capacity to neutralize multiple tier 2 strains, including T250.4, H703.2631, and CT184.D3.16 (Figures 3A-B, S2A-B).

**Figure 3.**
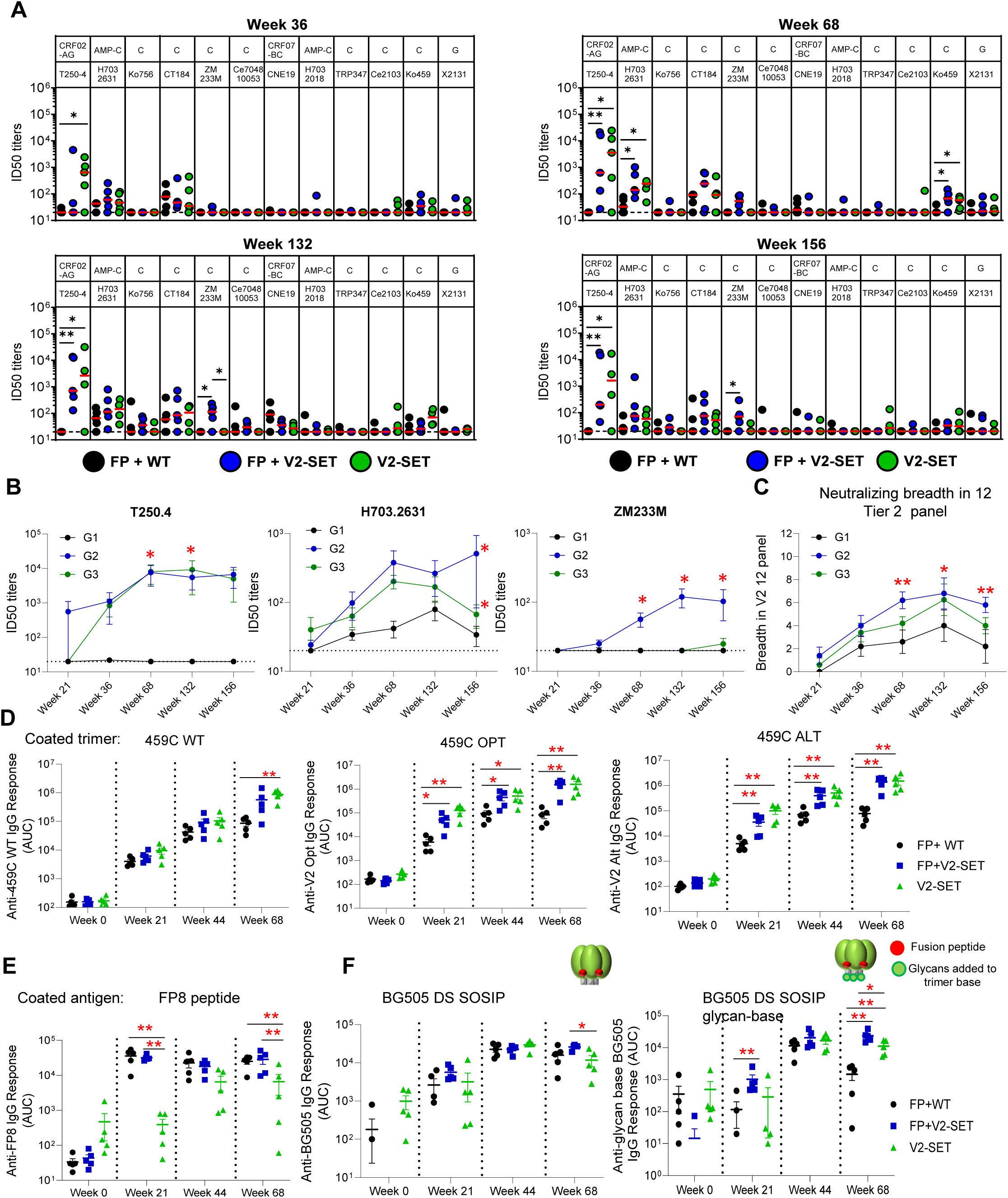
V2-SET trimer and FP-conjugate result in broader heterologous neutralization. (A) Serum neutralizing ID50 titers on 12 heterologous tier 2 virus panel at 4 different time points after immunization. Clade and virus IDs are listed on the upper and lower rows, respectively. (B) Kinetics of ID50 titers on T250.4, H703.2631 and CT184.D3.15 viruses. (C) The number of viruses neutralized within 12 virus panel in A. (D-F) Plasma response to vaccine-matched 459C WT, OPT and ALT trimers (D), FP peptide (E), and heterologous BG505 DS SOSIP trimer and its glycan-base trimer (F) at week0, week21, week 44 and week 68 are shown as the mean area under the curve (AUC) ± SEM. Two-tailed Mann-Whitney nonparametric tests were used for statistical analysis to estimate p values for different comparisons. *: p<0.05; **: p <0.01. See also Figure S2 and S3.

G2 differs from G3 by the addition of the FP8-rTTHC immunogen. To assess the impact of the FP8-rTTHC cocktail immunization, we compared G2 versus G3 sera for neutralizing activities and binding responses. We observed that for autologous neutralizing activity, FP8-rTTHC + V2-SET immunization in G2 exhibited slightly lower ID50 titers compared to V2-SET alone immunization in G3. However, for heterologous tier 2 virus neutralization, FP8-rTTHC + V2-SET elicited earlier and higher neutralizing activity against tier 2 viruses H703.2631, ZM233M (Figure 3B) and CT184.D3.15 (Figure S2B). Of note, only FP8-rTTHC + V2-SET immunized animals in G2 showed neutralizing activity on ZM233M (Figure 2B). Furthermore, FP8-rTTHC + V2-SET elicited broader neutralizing activity in the 12 tier 2 virus panel (Figures 3A-C, S3).

We next looked at binding antibody responses and observed that V2-SET immunized animals in G2 and G3 elicited significantly higher serum binding antibodies to vaccine-matched 459C WT, OPT and ALT trimers compared with 459C WT immunized animals in G1 (Figure 3D). As expected, FP8-rTTHC with WT trimer or with V2-SET trimers elicited significantly higher FP8-specific serum responses than did V2-SET alone, although V2-SET alone also elicited detectable FP-specific responses (Figure 3E). Interestingly, FP8-rTTHC addition elicited higher cross-strain responses to soluble BG505 DS SOSIP trimer, and the differences were more significant against a BG505 DS SOSIP glycan-base trimer (Duan et al., 2024; Olia et al., 2023), which has glycans added on the trimer base to block the base response, indicating the FP8-rTTHC addition increased the non-base antibody responses. As higher non-base responses have been correlated with higher neutralizing activity (Corrigan et al., 2021; Duan et al., 2024), we analyzed the correlation between serum response to BG505 DS SOSIP glycan base-trimer and neutralizing activity. We found a positive correlation between the week 68 serum binding antibody responses to BG505 glycan-base trimer and neutralizing titers at week 68 and week 132 against all three 459C pseudoviruses (Figure S2C-D).

### FP-specific mAbs from FP-rTTHC + V2-SET Env immunized animals demonstrate up to 47% neutralization breadth

We isolated FP-specific mAbs from all three vaccinated groups using FP and trimer probes as previously described (Kong et al., 2019; Kong et al., 2016; Xu et al., 2018). Utilizing PBMCs from week 68 after the first immunization series, FP-specific memory B cells were isolated using FP9-PEG12 and 459C V2-SET trimers as probes (Figure 4A). Antibody variable domains of the isolated B cells were sequenced, and selected clones were synthesized and expressed. As expected, isolated FP-specific mAbs could bind FP8 peptide and vaccine-matched 459C trimers (Figure S5A-C). Multiple mAbs exhibited cross-clade neutralizing activity, with two mAbs, J601-1A6 and J601-1B2, isolated from G2 FP+459C V2-SET immunized animal J601, neutralizing all 17 FP-sensitive strains tested, including human FP-specific bNAb VRC34.01-resistant strains CNE15 and 286.36 (Figure 4B).

**Figure 4.**
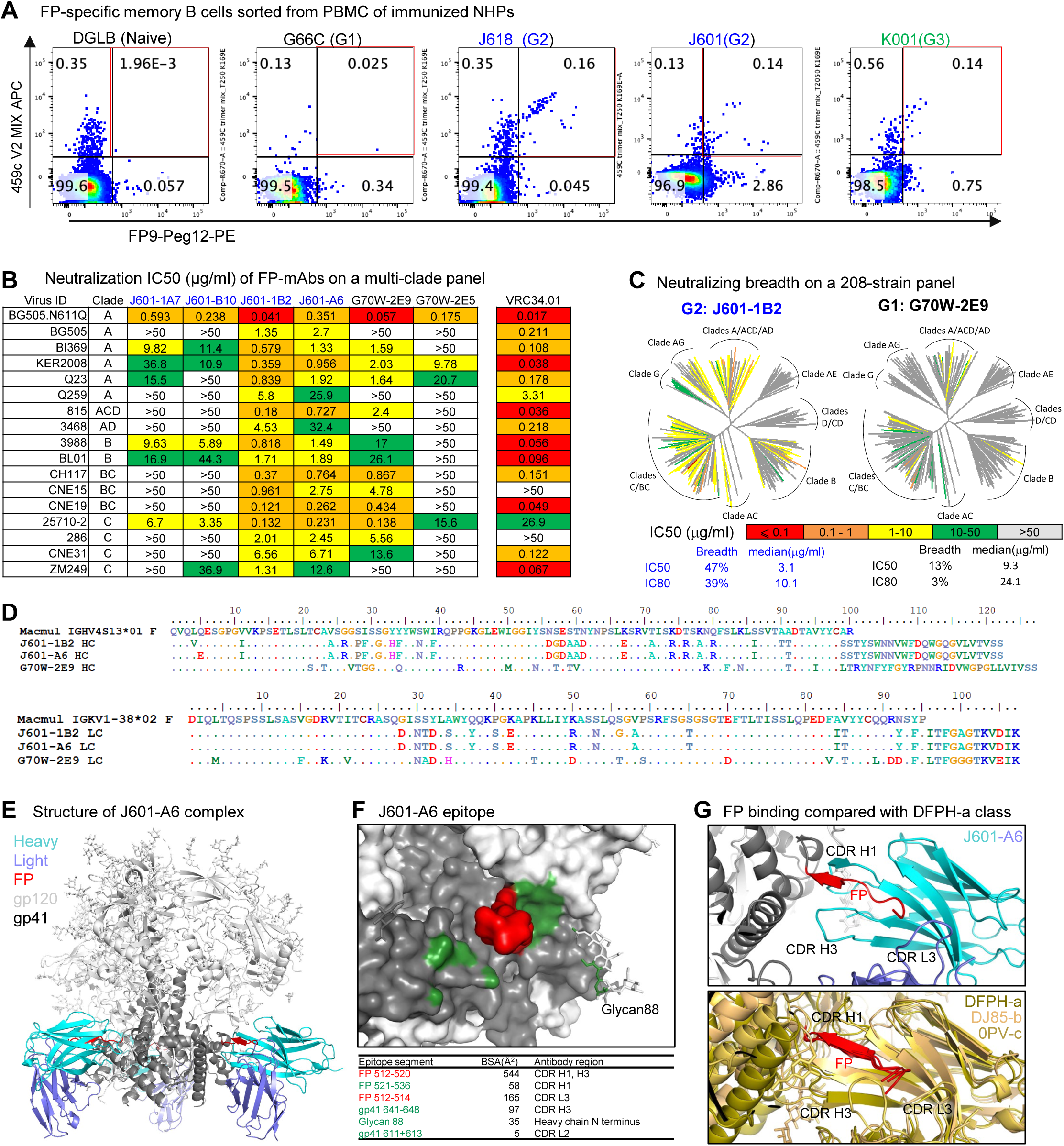
FP-specific mAbs isolated from FP+V2 SET immunized NHPs showed extensive maturation and up to 47% breadth on a 208-strain panel. (A) FP-specific mAbs were isolated with FP and trimer probes with FACS single-cell sorting. (B) FP-specific mAbs showed heterologous neutralization on FP-specific tier 1 and tier 2 panels, and VRC34 was used as a positive control for the panel and shown on the right column. (C) Two mAbs were tested on 208-strain panel, with neutralization breadth shown under the diagrams. (D) Genetic analysis of the two mAbs tested in 208-strain panel with an additional mAb J601-1A6 isolated from the same NHP. (E) Structure of J601-A6 in complex with Env trimer at 3.5 Å. J601-A6 binds at the FP site of vulnerability. (F) The J601-A6 epitope shown at one of the binding sites on the trimer, colored red for FP residues 512-520 and green for the rest of the epitope surface. Buried surface areas of the epitope segments are listed below the figure. The antibody binding is dominated by interacting with the N-terminal segment of FP. (G) FP binding interactions with J601-A6 (upper panel) compared with the broad DFPH-a class antibodies (lower panel), DFPH-a.15 (6N1W), DJ85-b.01 (8TKC), and 0PV-c.01 (PDB: 6NF2). The antibody heavy chains were aligned. The FP structures superposed well, but the rest of gp41 was shifted.

We tested J601-1B2 and another mAb G70W-2E9, isolated from G1 FP + 459C WT immunized animal G70W, on a 208-strain HIV-1 panel. J601-1B2 neutralized 47% of tested viruses with an IC50 <50 μg/ml and neutralized 39% of tested viruses with an IC80 <50 μg/ml. G70W-2E9 neutralized 13% and 3% of viruses with an IC50 <50 μg/ml and IC80 <50 μg/ml, respectively. In addition, J601-1B2 was more potent than G70W-2E9 (median IC50 3.1 μg/ml vs 9.3 μg/ml, median IC80 10 μg/ml vs 24 μg/ml), although both were less potent than the human FP-specific bNAb VRC34.01 (Figure S4A-B). Of note, all neutralizing FP-specific mAbs were isolated from FP8-rTTHC immunized G1 and G2 animals. FP-specific mAbs were also isolated from G3 animals immunized with V2-SET trimer alone, but these mAbs were not neutralizing (Figure S4A). As 459C V2-SET trimers were used in G2, we tested the binding of FP-neutralizing mAbs to 459C WT, OPT, and ALT trimers, and all these mAbs bound to 459C WT slightly less than to OPT and ALT trimers (Figure S5A-C).

### FP-specific mAbs bind to Env trimer with binding modes similar to the previously identified DFPH-a class FP-specific bNAbs

To elucidate the structural interactions of the isolated FP-specific mAbs with Env, we determined cryo-EM structures of J601-A6 and J601-1B2 Fabs, each in complex with the BG505 DS-SOSIP Env trimer. J601-A6 and J601-1B2 were derived from the same lineage, with differences in only three residues in the heavy chain and one in the light chain (Figure 4D). The cryo-EM structure of J601-A6 was resolved at 3.5 Å and that of J601-1B2 at 3.8 Å (Figure 4E, Data S1.4, S1.5, S1.8). The two structures are essentially identical, with the distinguishing residues away from the antibody interface with the Env trimer (Data S1.5G, H). Therefore, the effect of these mutations on binding and neutralization is indirect and subtle. J601-A6 binds at the FP site, interacting primarily with the FP N-terminal residues 512-520 (Figure 4E-G). The dominance of binding to FP N-terminal segment allows flexibility in the antibody approach angles, resulting in multiple 3D classes of cryo-EM particles with slightly different binding orientations of the antibodies on the trimer (Data S1.4C, D; Data S1.5E, F). The FP N-terminal segment is held between the complementarity determining region 3 of the heavy chain (CDRH3) and CDRH1 and interacts with CDRL3 (Figure 4G). The binding mode of J601-A6 and J601-1B2 is similar to the broadest vaccine-elicited DFPH-a class FP bNAbs (Kong et al., 2019; Wang et al., 2024) (Figure 4G; Data S1.4G, H). The dominant binding to the relatively conserved FP N-terminal segment allows for the broad recognition of HIV-1 strains, although the moderate binding surface area and low somatic hypermutation appear to limit the potency of neutralization.

### V2-SET Env immunized G2 and G3 animals show V2-specific serum neutralization

As we observed higher autologous and broader heterologous neutralization with the V2-SET trimers, we tested sera for neutralization against V2 epitope mutants N160A and K169E on a T250-4 Env backbone, with the V2 apex-specific bNAbs PG9 and CAP256/VRC26.25 included as positive controls and the V3-glycan bNAb PGT121 as a negative control. As expected, PGT121 was not impacted by N160A or K169E mutations, whereas V2 apex-specific bNAbs lost neutralization activity on N160A and K169E mutants, except CAP256/VRC26.25, which retained strong neutralization on N160A. Week 68 sera from V2-SET immunized G2 and G3 animals were tested, and all showed substantial reductions of neutralization to the K169E mutant, indicating that serum neutralization against T250-4 was due to V2-specific NAbs. However, all animals except K120 showed enhanced neutralization on the N160A mutant, indicating that this mutation is beneficial for serum neutralization (Figure 5A, B). We also assessed serum neutralization against ZM233.6 and similarly found that animals from V2-SET immunized G2 showed reduction of neutralization to the K169E mutant, indicating that serum neutralization against ZM233.6 was also due to V2-specific NAbs (Figure 5C, D). These data demonstrate that V2-specific NAbs contributed substantially to the serum neutralization.

**Figure 5.**
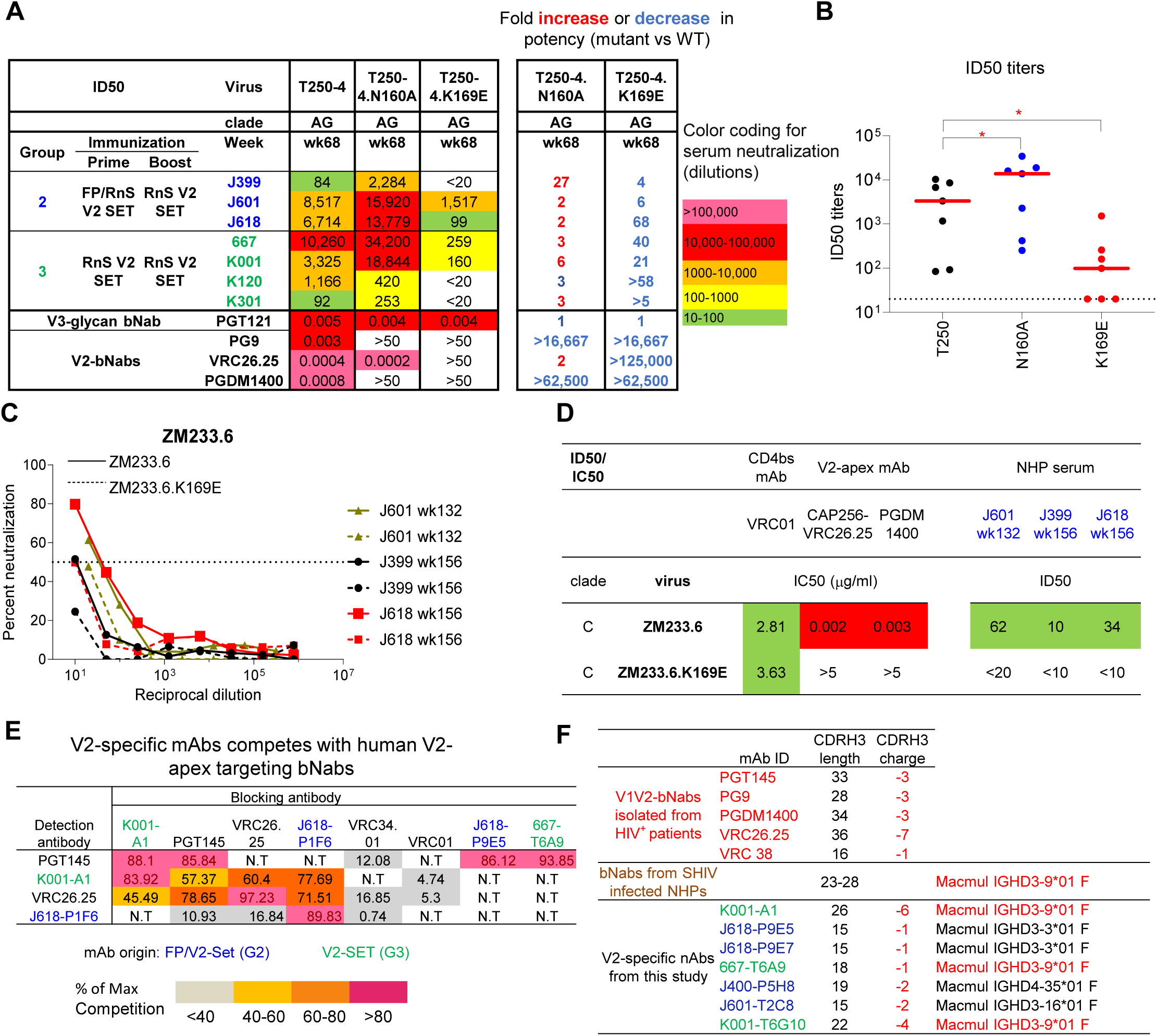
V2-SET trimer immunized NHPs showed V2-specific serum neutralization, with isolated mAbs showing genetic signatures/binding modes similar to previously identified FP and V2-specific bNAbs. (A) V2-SET immunized G2 and G3 NHPs with T250.4 neutralizing activity were tested on T250.4 V2-mutants N160A and K169E. ID50 fold change relative to WT T250.4 were shown on the right column. (B) ID50 titers from A are shown in scatter dot plots with lines representing median ID50 for each virus. (C) Percent of serum neutralization at serial dilutions on ZM233 and its K169E mutant. (D) ID50 titers of NHP sera on ZM233 and its K169E mutant as shown in (C). CD4bs-bNab VRC01, V2-apex bNAbs CAP256/VRC26.25 and PGDM1400 used as QC for the assay. (E) V2-specific mAbs competed with human V2-specific bNabs for soluble trimer binding. (F) Genetic signatures of V2-specific bNabs isolated from indivuals infected with HIV-1, SHIV infected NHPs and V2-SET immunized NHPs from this study. Statistical analysis was performed with the 2-stage linear step-up procedure of Benjamini, Krieger and Yekutieli to assess p-values for different groups. *: p<0.05; **: p <0.01. See also Figure S4, and Data S1.

### V2-specific mAbs isolated from V2-SET Env immunized animals showed neutralization with binding modes similar to the previously identified V2 bNAbs

To further investigate the serum antibody neutralization activity observed in 459C V2-SET immunized G2 and G3 animals, we used 459C WT, OPT, ALT, T250.4, ZM233.6, H703.2631 and CT184 gp140 DS SOSIP trimer probes and K169E mutant probes for V2-specific mAb isolation (Figure S5A). We identified V2 apex-targeted mAbs that effectively competed with V2 apex bNAbs on binding to Env trimers (Figures 5E, S5B). Antibody sequence analysis revealed well-characterized V2 bNAb signatures, including a long and negatively charged CDRH3 and the usage of the Mamu-IGHD3-9*01 gene (Figure 5F).

Next, we tested the V2-specific mAbs in neutralization assays and classified them based on the neutralization pattern of 459C WT, OPT, and ALT pseudoviruses. Neutralization class I mAbs neutralized 459C OPT but not WT and ALT pseudoviruses; neutralization class II mAbs neutralized 459C WT and ALT but not OPT pseudoviruses; and neutralization class III mAbs neutralized all three autologous pseudoviruses and some heterologous pseudoviruses, including T250.4, ZM233, and Ko459 (Figure 6A). CDRH3 sequence alignment of three classes of mAbs showed similar motifs (Figure 6B). In binding assays, these antibodies showed stronger binding to 459C OPT, ALT than WT trimers (Figure S5C). All T250.4 neutralizing mAbs showed reduced activity against N160A and K169E mutants in neutralization assays, confirming that these mAbs were V2 apex-specific mAbs. Indeed, negative-stain EM 3D reconstruction maps of several of these antibodies revealed their binding to the V2 apex of Env trimers (Figure 6D). Comparison of the negative-stain EM 3D map of the K001-A1 Env trimer complex with the cryo-EM map of PG9 Env trimer complex (EMD-25736) (Willis et al., 2022) indicated a similar binding mode but with a slightly different angle of approach (Figure 6E).

**Figure 6.**
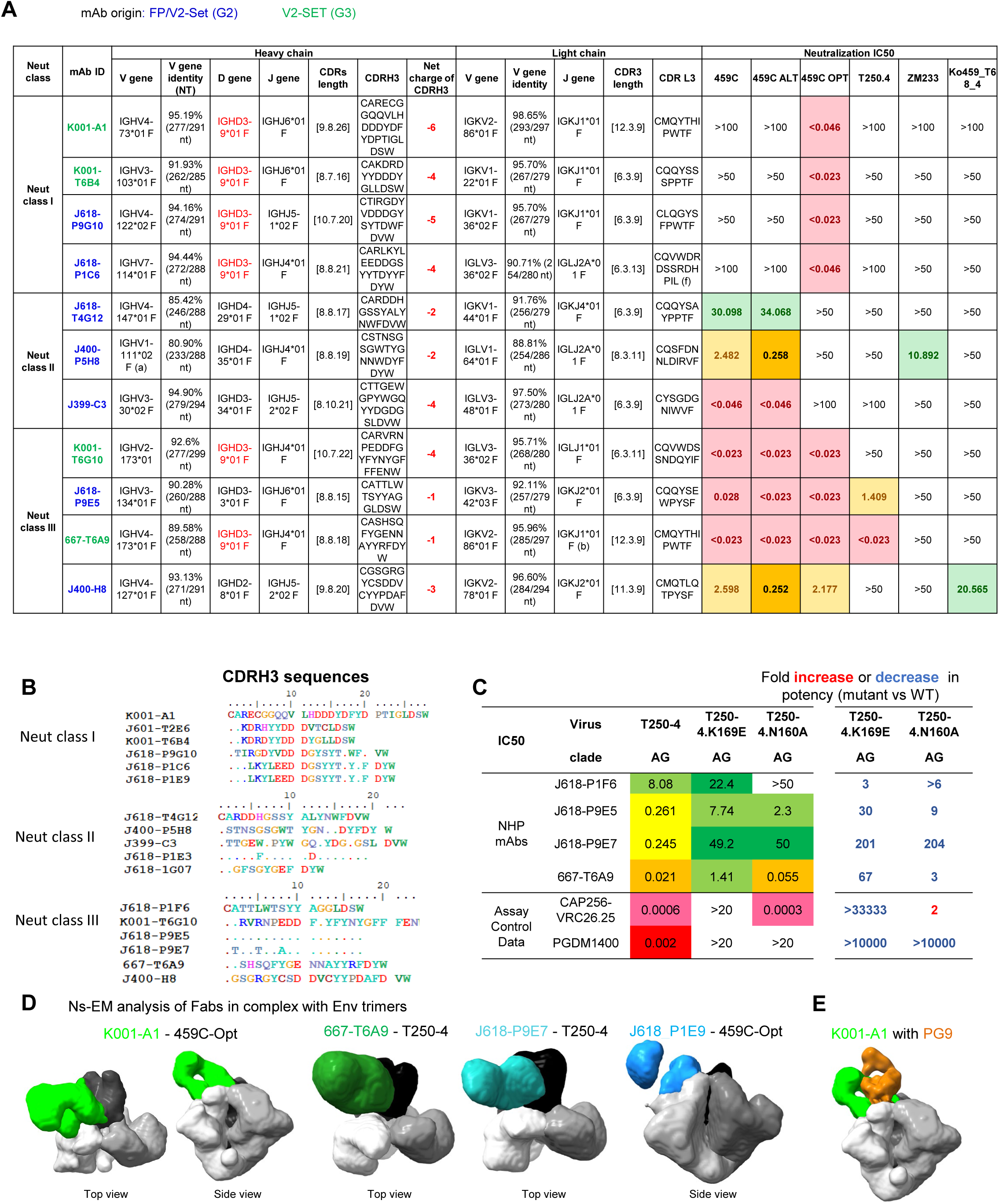
V2-specific mAbs isolated from V2-SET immunized G2 and G3 NHPs showed autologous and heterologous neutralization on tier 2 viruses, with V2-specific neutralizing signature and binding modes. (A) Three classes of V2-specific mabs were isolated and grouped based on neutralizing activity on 459C WT, OPT and ALT viruses. mAbs isolated from Group 2 and 3 are labeled blue and green fonts, respectively. IGHD3-9*01 is labeled in red. (B) CDRH3 amino acid sequence alignment of the three class V2-specific antibodies shown in (A). (C) mAb neutralization on T250.4 is abolished with K169E mutant. ID50 fold change relative to WT T250 are shown on the right columns. (D) V2-specific mabs bind to V2-apex on Env trimer by nsEM analysis. (E) Superposition of nsEM map of K001-A1 complex with cryo-EM map of PG9-Env complex (EMD-25736). See also Data S1.

### K001-A1 binds the V2 apex of the Env trimer, interacting with all three protomers at the V2 apex hole and tilting to contact primarily the C-strand of one protomer

To characterize the molecular interactions between K001-A1 mAb and Env trimer, we determined a cryo-EM structure of K001-A1 Fab in complex with 459C-OPT Env trimer at 3.0 Å (Figure 7, Data S1.6-1.8). There is a single copy of K001-A1 Fab binding at the V2 apex of the Env trimer interacting with the three gp120 subunits around the apex hole (Figure 7A, Data S1.7A). The cryo-EM map has sufficient density for the K001-A1 constant domains, albeit weaker than the rest (Data S1.7E), which enabled modeling the Fab constant domains. The antibody binding mode is analogous to PG9 (Davenport et al., 2011) and CAP256/VRC26.25 (Gorman et al., 2020) with the CDRH3 inserting into the apex hole and the rest of the binding epitope predominantly but not exclusively on one gp120 subunit.

**Figure 7.**
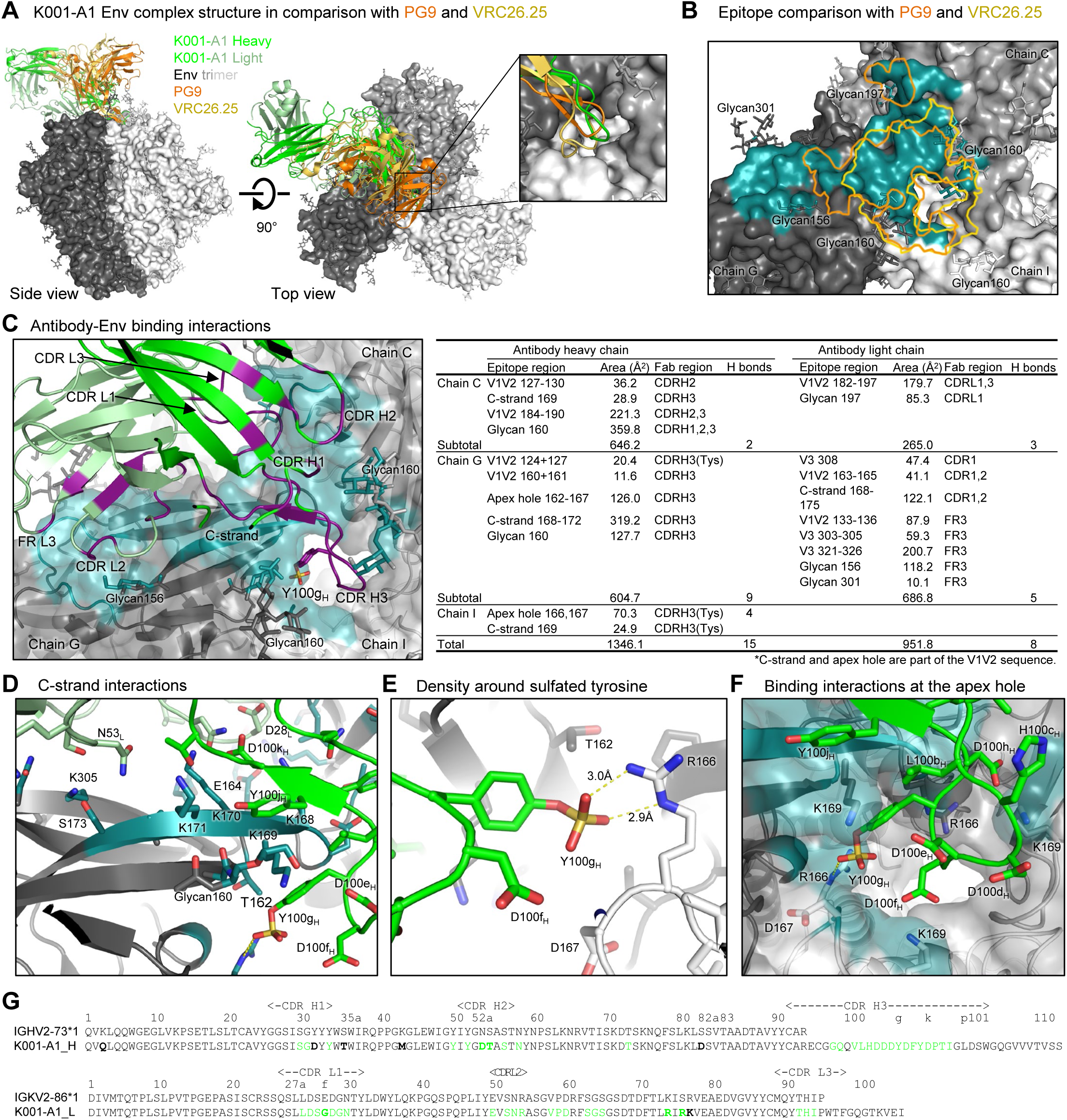
Structure reveals antibody K001-A1 to bind both V2 apex and sequence-variable regions, suggesting its maturation was inhibited because it binds only OPT. (A) Cryo-EM structure of K001-A1 Fab in complex with 459C-OPT-RnS Env trimer at 3.0 Å, superimposed with the PG9 and CAP256/VRC26.25 complexes. The structures were aligned by the Env trimers, with only the trimer of the K001-A1 complex shown as gray surface with glycans as sticks. The inset is a zoom-in at the apex hole. (B) Zoom in on the K001-A1 epitope with superimposed outlines of PG9 and CAP256/VRC26.25 epitopes aligned based on the Env trimer structures. The K001-A1 epitope is colored teal for the protein surface and glycan atoms. The three gp120 subunits in the trimer are designated as chains G, C, and I, as labeled. (C) K001-A1 binding interactions with Env trimer. Paratope residues are highlighted in purple; epitope residues are highlighted in teal. The buried surface areas of the epitope regions were calculated with PISA (https://www.ebi.ac.uk/pdbe/pisa/). (D) Zoom in on the C-strand to reveal details of Fab binding interactions. (E) The sulfated tyrosine interacted with the charged side chain of R166. (F) Details of the binding interactions around the apex hole. (G) K001-A1 heavy and light chain sequences, aligned with rhesus monkey germline variable sequences, with Kabat numbering. Somatic hypermutations are indicated by bold fonts; residues in contact with Env trimer are indicated by green fonts. See also Data S1.

K001-A1 binding differs from PG9 and CAP256/VRC26.25 in the relative position of heavy and light chains and a larger tilt angle, and its light chain contacts the trimer. Consequently, the K001-A1 epitope area is larger with an additional patch distal to the apex hole, unlike PG9 and CAP256/VRC26.25 (Figure 7B). CAP256/VRC26.25 has its CDRH3 loop penetrating deeply into the apex hole and thus interacts most strongly with all three gp120 subunits around the apex hole. K001-A1 is more like PG9, with a rather asymmetric epitope surface, interacting with two of the three glycans at 160 and reaching to glycan 197 and glycan 156. The interactions with glycan 156, however, are different between PG9 and K001-A1 (Data S1.7C). PG9 interacts with glycan 156 favorably, maintaining its ligand-free conformation. K001-A1, however, would clash with glycan 156 if this glycan adopts the same conformation as in the PG9 complex or the glycan 156 of the other two protomers in the K001-A1 complex; instead, the K001-A1 light chain pushes away glycan 156, which is mostly disordered in the cryo-EM density map. Therefore, K001-A1 interaction with glycan 156 is likely energetically unfavorable.

K001-A1 binding to the Env trimer involves both heavy and light chains. The heavy chain interactions are similar to other V2 apex bNAbs, involving all three CDRs binding the V1V2 region, with the long CDRH3 dominating the binding (Figure 7C). The heavy chain contact regions are around the apex hole, involving mostly chains C and G (the three gp120 subunits were designated as chains C, G, and I, with chain G containing most of the binding epitope; Figure 7B, C). The K001-A1 light chain interactions involve all three CDRs and the framework region 3 (FR L3), interacting with mostly chain G and covering an additional patch away from the apex hole. The total epitope surface area is nearly 2300 Å^2^, which is larger than that of PG9 and CAP256/VRC26.25 (Data S1.7D, E).

Despite the substantial contribution to the epitope from the light chain binding, the heavy chain binding mimics the binding of the well-characterized V2 bNAbs, with its long, highly negatively charged CDRH3 interacting with the positively charged residues at the apex hole and the lysine-rich V1V2 β-strand C (C-strand) (Figure 7C-G). The C-strand plays a dominant role in binding interactions of V2 apex bNAbs (Gorman et al., 2020; Roark et al., 2025; Willis et al., 2022) 459C OPT Env has four lysine residues at positions 168-171. In chain G, the gp120 subunit contributes to the majority of the epitope, and all the lysine side chains are in close contact with K001-A1 (Figures 7D). K168 is 5.2 Å away from D28_L_; K169 is 4.6 Å from D100e_H_; K170 is 2.8 Å from D100k; and K171 is 3.2 Å from the carbonyl of P100l_H_ and has hydrophobic interactions with P100l_H_ and Y100j_H_. Most of these lysine residues within the epitope are variable among 459C WT, OPT, and ALT trimers, as well as among the circulating HIV-1 strains (Data S1.7B, S1.9), partially explaining the limited heterologous neutralization by K001-A1; the variability in Env positions 168-171 also generally constrains the breadth and potency of V2 bNAbs (Bricault et al., 2019). Moreover, Y100 of the CDRH3 is sulfated and has a salt bridge with R166 of chain G (Figure 7E). This sulfated tyrosine adds to the highly negative charges of the CDRH3, which interacts with the positively charged R166 and K169 residues at the apex hole from the three protomers (Figure 7F). Overall, the K001-A1 binding has substantial similarities to PG9, with parallel β-strand and charge pairing with the C-strand and the apex hole, as well as the axe recognition mode.

## DISCUSSION

Elicitation of bNAbs by vaccination has been a long-sought goal for the HIV-1 vaccine field, and induction of bNAbs to multiple sites of vulnerability will almost certainly be required for an effective HIV-1 vaccine given the challenge of HIV-1 variability. However, most HIV-1 vaccine studies have to date focused on the induction of bNAbs to a single epitope, such as the FP, V2 apex, V3 supersite, and CD4 binding site (Martinez-Murillo et al., 2017; Mogus et al., 2020; Voss et al., 2017; Wang et al., 2024; Xu et al., 2018). In this study, we designed a combined dual-vaccine and demonstrated in rhesus macaques simultaneous induction of NAbs to both V2 apex and FP, two key targets for HIV-1 vaccine development.

Our dual-vaccine included a FP-rTTHC conjugate and three related 459C V2-SET Env trimers with V2 modifications (Bricault et al., 2019; Xu et al., 2018). The FP-rTTHC immunogen was co-administered with trimers at the start of the vaccination regimen (Cheng et al., 2020; Corrigan et al., 2021), and the trimers had compatible FP sequences. The 459C OPT trimer had enhanced binding to V2-specific bNAbs, longer retention in draining lymph nodes, and the ability to induce autologous neutralizing titers >1000-fold higher than the 459C WT trimer. Vaccine-elicited autologous tier 2 NAb titers reaching >1,000,000 are uncommon for neutralization-resistant HIV-1 strains such as 459C. These NAb titers contrast with the relatively modest autologous neutralizing titers elicited by 459C WT trimer, which were low at both weeks 68 and 132 (a single animal reached a maximum NAb titer of 323). The altered immunogenicity was not apparent antigenically (Fig. 1B), but differences were observed in the EM analysis (Data S1), with the wildtype 459C trimer producing well-ordered reconstruction at 3 Å resolution, while the OPT trimer was too flexible to enable residue-level tracing. It remains to be seen whether the enhanced lymph node retention and the increased Env immunogenicity using this strategy can be translated to other Env strains.

In addition to autologous and heterologous tier 2 serum NAb responses, both V2 apex-specific and FP-specific mAbs were isolated from the immunized animals and showed both autologous and some heterologous neutralizing activity. In addition, V2 apex-specific mAbs showed competition against human V2-specific bNAbs on binding to Env trimers, sequence analyses revealed similar antibody signatures, including long and negatively charged CDRH3, and structural analyses revealed V2 apex binding of these isolated mAbs. However, V2-specific mAbs showed only limited heterologous neutralizing breadth, likely because of limited maturation and increased specific light chain contacts. These data suggest a path forward for improving induction of V2-specific NAb breadth by adding immunogens that facilitate antibody maturation. In contrast, FP-specific mAbs showed broad cross-strain neutralizing activity with 47% breadth on a 208-strain HIV-1 panel.

In summary, our data demonstrate that FP-rTTHC conjugate and 459C V2-SET Env immunization elicited NAbs against both FP and V2 apex epitopes that exhibited potent autologous neutralizing activity and some cross-strain heterologous neutralizing breadth in rhesus macaques. Although most HIV-1 vaccine studies have to date focused on induction of NAbs against a single epitope, the strategy to target multiple sites of HIV-1 Env vulnerability simultaneously will likely be critical for an effective HIV-1 vaccine for humans.

### Limitations of Study

Our analyses of epitope-specific antibody responses involve macaques immunized with FP8-rTTHC and 459C Env trimers, so it is possible that immunization with different FP peptides, Env trimers, base region sequences, and aduvants may show different results. In addition, the addition of an antigen that improves maturation of the V2 NAb responses will likely be required to improve neutralization breadth. Finally, clinical trials will be required to demonstrate the translatability of our findings for humans.

### Consortia

The VRC Production Program includes N. Amharref, F.J. Arnold, P. Bandi, N. Barefoot, C. Barry, E. Carey, R. Caringal, K. Carlton, N. Chalamalsetty, A. Charlton, R. Chaudhuri, M. Chen, P. Chen, Y. Chen, N. Cibelli, J.W. Cooper, H. Dahodwala, G. Dobrescu, M. Fleischman, J.C. Frederick, H.C. Fuller, J.G. Gall, I. Godfroy, D. Gollapudi, D. Gowetski, K. Gulla, J. Horwitz, A. Hussain, V. Ivleva, T. Khin, L. Kueltzo, G. Lagos, Q. Paula Lei, Y. Li, R. Luedtke, V. Mangalampalli, G. Moxey, S. O’Connell, A. Patel, E. Rosales-Zavala, E. Scheideman, N.A. Schneck, Z. Schneiderman, A. Shaddeau, W. Shadrick, S. Shetty, R. Skubutyte, B. Tippett, A. Vinitsky, C. Webber, S. Witter, L. Yang, Y. Yang, and Y. Zhang.

## Acknowledgments

We thank members of the Virology Laboratory, Vaccine Research Center, for discussions and comments on the manuscript. We thank D. Ambrozak for helping with FACS sorting. We thank R. Carroll, N. Jean-Baptiste, C. Moore, C. Whittaker, and A.B. McDermott for their assistance with neutralization assessments on the 208-strain panel, and J. Baalwa, D. Ellenberger, F. Gao, B. Hahn, K. Hong, J. Kim, F. McCutchan, D. Montefiori, L. Morris, E. Sanders-Buell, G. Shaw, R. Swanstrom, M. Thomson, S. Tovanabutra, C. Williamson, and L. Zhang for contributing HIV-1 envelope plasmids used in neutralization. We thank the VRC Production Program for providing the FP-conjugate immunogen. This research was supported in part by the Intramural Research Program of the National Institutes of Health (NIH). The contributions of the NIH authors are considered Works of the United States Government. The findings and conclusions presented in this paper are those of the authors and do not necessarily reflect the views of the NIH or the U.S. Department of Health and Human Services. This research was supported in part by the International AIDS Vaccine Initiative’s (IAVI’s) Neutralizing Antibody Consortium. The animal study was supported by Gates Foundation grants INV-048656 (D.H.B.). Some of the neuralization assays were supported by Gates Foundation INV-036842 (M.S.S.). Support for this work was also provided by NIH grants AI164556, AI169615, and AI177687 (D.H.B.). Part of the cryo-EM data collection was performed at the Columbia University Cryo-Electron Microscopy Center, which was funded in part by the Frederick National Laboratory for Cancer Research, NIH, under contract HHSN261200800001. We also thank the NIAID Research Technologies Branch, Microscopy Unit for Cryo-EM for data collection. Support of this work was also provided by Leidos Biomedical Research contract S23-012 (L.S., Z.S.). This study used the NIAID Office of Cyber Infrastructure and Computational Biology High Performance Computing cluster.

## Author Contributions

DHB and PDK led the overall study. HD led the mAb isolation and characterization, figure preparation and manuscript drafting; HD, JPN, SW and JC drafted the manuscript; HD and JPN led the serum analysis and figure preparation; JPN prepared the immunization and sample collection; SW performed structural analysis for the isolated mAbs; JC, AR NL and AEP performed mouse study and data analysis; ITT made biotinylated trimer probes for the B cell sorting and FACS analysis, performed BLI antigenicity analysis; HD, GSS, DKP and ASC performed the single-cell nested PCR and sequencing; BZ helped HD with mAb plasmid synthesis; HD, DKP, MC, AB and ASC performed mAb transfection and purification; ARH, CS, SK helped with single-cell RATP-ig analysis; SOD, MK, HW and CL performed neutralization and data analysis; HD and CHS performed sequence analysis for mAbs; HL and LR helped with immunogen preparation and characterization; CC supervised GSS, DKP, AB and ASC; YY helped HD with mAb transfection; JEB, NCM, and RSR collected cryo-EM data and SW carried out structural analysis: BZ, HD, DJVW made mAb Fab preparations for EM analysis;RR helped with plasmid design; RC, MM performed 208 virus panel neutralization for mAbs; NADR and LW supervised SOD and MK for neutralization; DCD supervised ARH, CAS supervised SK for RATP-ig analysis; RAK, JRM and TCP supervised HD; BK helped with immunogen design; PDK, LS and TZ supervised SW, ITT, BZ, YY, NCM, HD, DVW, ASO, JEB, RSR; MSS supervised CL; TCP reviewed data, mentored HD; PDK and DHB initiated the animal immunization; TCP, PDK and DHB headed the serum analysis and mAb isolation and characterization, reviewed the data and figures. All authors provided comments and revisions to the manuscript.

## RESOURCE AVAILABILITY

### Lead Contact

Further information and requests for resources and reagents should be directed to and will be fulfilled by Theodore C. Pierson, Peter D. Kwong and Dan H. Barouch.

### Materials Availability

All new reagents are available by MTA for non-commercial research.

### Data and Code Availability

The published article includes all data generated or analyzed during this study. This study did not generate new code.

## EXPERIMENTAL MODEL AND SUBJECT DETAILS

### Macaque Study

15 outbred rhesus macaques of Indian origin (*Macaca mulatta*) were assigned to three study groups (n=5/group) with even sex and weight distributions. All animals were housed and maintained at Alpha Genesis Inc. (Yemasee, SC) which is fully accredited by the Association for Assessment and Accreditation of Laboratory Animal Care International (AAALAC) and approved by the Office of Laboratory Animal Welfare (NIH/PHS Assurance ID: D16-00387). All procedures were conducted in compliance with all relevant local, state, and federal regulations and were approved by the Alpha Genesis Institutional Animal Care and Use Committee (IACUC). Group 1 received three bimonthly immunizations of rTTHC.FP8.v1 (100µg) mixed with 459C repair and stabilized (RnS) wild-type (WT) gp140 Env (100µg) at weeks 0, 8 and 16 followed by six bi-monthly immunizations of 459C RnS WT gp140 Env at weeks 24, 32, 40, 48, 56 and 64. After a resting interval of 48 weeks, animals received an additional five bimonthly booster immunizations of rTTHC.FP8.v1 (100µg) combined with 459C RnS WT gp140 Env (100µg) at weeks 112, 120, 128, 136, 142 with a sixth and final boosting immunization of 459C RnS WT gp140 only at week 152. Group 2 received the same immunization regimen as group 1 with 459C RnS WT gp140 Env replaced with trivalent 459C V2-SET gp140 Env (100µg total). Group 3 received the same regimen as groups 1 and 2 using trivalent 459C V2-SET (100µg total) only. All immunizations were administered intramuscularly and bilaterally in quadriceps muscles using Adjuplex adjuvant.

### Antigen labeling and characterization

HIV Env trimer protein antigens (WT, OPT, and ALT) were conjugated to Alexa Fluor 647 (A647) using the Alexa Fluor 647 Protein Labeling Kit (Invitrogen, A20173), following the manufacturer’s protocol. Env trimers were first diluted to 2 mg/mL in 0.5 mL of PBS. To this, 50 µL of 1 M sodium bicarbonate buffer (pH 8.4) was added, and the mixture was maintained on ice. The buffered protein solution was then added to the vial containing the A647 dye. Labeling reactions were carried out at RT for 2 hours with gentle stirring. Following conjugation, excess dye was removed by passing the samples through Zeba Spin Desalting Columns pre-equilibrated with PBS, with two rounds of purification. Labeled proteins were sterile-filtered using 0.22 µm Spin-X centrifuge tube filters (Millipore Sigma, CLS8160) and stored at 4°C until further use. The degree of labeling was determined by measuring absorbance at 280 nm and 650 nm, corresponding to total protein and A647 dye, respectively. Protein and dye concentrations were calculated using extinction coefficients of 118,000 M⁻¹cm⁻¹ for each trimer subunit (WT, OPT, and ALT) and 239,000 M⁻¹cm⁻¹ for the A647-NHS ester. The labeling ratio was calculated as the molar ratio of A647 dye to trimer protein.

### Mice immunization and sample collections

All animal procedures were approved by the Institutional Animal Care and Use Committee (IACUC) of Beth Israel Deaconess Medical Center (BIDMC) and conducted in compliance with the NIH Guide for the Care and Use of Laboratory Animals, the Animal Welfare Act of the U.S. Department of Agriculture, and in an AAALAC-accredited facility, with IACUC protocol number 050-2022. A total of 72 eight-week-old female Balb/cJ mice (strain #000651, The Jackson Laboratory) were randomly assigned to four experimental groups (n=18). Mice were housed in pathogen-free conditions on a 12-hour light/dark cycle at 21 ± 1°C with ad libitum access to food and water. Mice were anesthetized and immunized intramuscularly with 50 μg of Alexa Fluor 647-conjugated immunogens (WT-A647, OPT-A647, or ALT-A647) formulated in a 1:1 ratio with Adju-Phos adjuvant (InvivoGen, vac-phos-250), administered in a total volume of 100 μL (50 μL per hind leg). A group of unimmunized animals served as naïve controls. Draining lymph nodes (dLNs) and spleens were harvested at days 1, 3, 5, 7, 9, and 12 post-immunization following euthanasia and placed into RPMI-1640 medium supplemented with 2% FBS for further processing.

### Flow cytometry analysis of draining lymph nodes and spleen

Following tissue collection, dLNs were enzymatically digested in 1 mL of digestion buffer containing 0.8 U/mL Dispase (STEMCELL), 0.1 mg/mL Collagenase D (Worthington Biochemicals), and 0.1 mg/mL DNase I (Sigma) at 37 °C for 20 minutes. Tissues were subsequently dissociated into single-cell suspensions by mechanical disruption through a 70-μm cell strainer (BD Biosciences). Spleens were processed without enzymatic digestion by mechanical disruption through a 70-μm filter, followed by red blood cell lysis using ACK lysis (Gibco) buffer for 3 minutes at RT. Single-cell suspensions from dLNs and spleens were washed with PBS and stained with Live/Dead Fixable Aqua Dead Cell Stain (Thermo Fisher Scientific, L34957) for 10 minutes at 25 °C. After two washes with MACS buffer supplemented with 2% BSA (Miltenyi) and 1.5% penicillin-Streptomycin (Fisher Scientific), cells were blocked with anti-mouse CD16/CD32 Fc receptor antibody (BioLegend, 101319) for 10 minutes at 4 °C, followed by surface staining with fluorophore-conjugated antibodies for 30 minutes at 4 °C. Flow cytometric analysis was performed on a BD LSR II flow cytometer (BD Biosciences). To identify immune cell populations associated with Alexa Fluor 647-labeled immunogens, the cells were stained with the CD3 BUV737 (clone 17A2, BD Biosciences), CD21/CD35 PE (clone 7E9, BioLegend), CD169 BV605 (clone 3D6.112, BioLegend), and B220 BUV395 (clone RA3-6B2, BD Biosciences).

### Mannose-Binding Lectin (MBL) deposition ELISA

High-binding 96-well ELISA plates were coated overnight at 4°C with WT, OPT, or ALT SOSIP protein at 1μg/mL diluted in 1X DPBS. After incubation, plates were washed once with wash buffer (0.05% Tween-20 in 1X DPBS) and then blocked with 350 μL/well of a casein-based blocking solution for 2–3 hours at room temperature (RT). After incubation, the blocking solution was discarded, and the plates were gently blotted dry. Serum was freshly collected from naïve BALB/c mice using Sarstedt serum gel tubes and kept on ice. Initial serum dilutions were prepared at 30% (v/v) in blocking buffer and further serially diluted two-fold. Diluted samples were added to the antigen-coated wells and incubated for 1 hour at 37°C. Following incubation, plates were washed three times with wash buffer and incubated with 2 μg/mL of rat anti-mouse MBL monoclonal antibody (clone 14D12, Abcam, cat# ab106046) prepared in blocking buffer and incubated for 1h at RT. After three washes, HRP-conjugated anti-rat IgG secondary antibody (1:5000 dilution; Invitrogen, cat# 31470) was added, and plates were incubated for 1h at RT. After three washes, 100 μL of SeraCare KPL TMB SureBlue substrate was added to each well and incubated for 11 minutes at RT. The reaction was stopped by adding 100 μL of SeraCare KPL TMB stop solution. Absorbance was measured at 450 nm with a reference reading at 650 nm using a VersaMax microplate reader (Molecular Devices). Endpoint titers were determined by fitting the data to a four-parameter logistic (4PL) regression model. The reciprocal serum dilution corresponding to a corrected absorbance value (A450–A650) of 0.1 was reported as the endpoint titer.

### Adjuvants

For each immunization, 200 μl of Adjuplex (Sigma-Aldrich Inc, MO or Adjuplex equivalent formulated based on US Patent 6,676,958 B2) was mixed with immunogen for adjuvant studies.

### Cell Lines

Expi293F cells (cat# A14257) and FreeStyle 293-F cells (cat# R79007) were purchased from ThermoFisher Scientific Inc. Cells were maintained in FreeStyle 293 Expression Medium. The cell line was used directly from commercial sources and cultured following the manufacturer’s suggestions, as described in the Method Details section below.

## METHOD DETAILS

### Expression of 459C V2-SET gp140 Env Constructs

The 459 V2-SET gp140 Env proteins were generated as previously described (Rutten L et al, Cell Rep. 2018, 23(2):584-595). Constructs were synthesized and codon-optimized (GenScript) and cloned into pCDNA2004 or generated by standard methods involving site-directed mutagenesis and PCR. HEK-Expi293 cells were transiently transfected with 90% of pCDNA2004 plasmid with the Env insert and 10% of Furin-pCDNA2004, according to the manufacturer’s instructions and cultured for 5 days at 37°C and 10% CO2. Culture supernatants were spun for 10 min at 1,250 × g. For expressions in 96-well format, the cells were cultured for 3 days at 37°C and 10% CO2. 4 μL of Opti-MEM was mixed with 4 μL 100 ng/μL DNA, and 8 μL Expi293F mix (54 μL/mL Opti-MEM) as added and incubated for 20 min. Subsequently, 200 μL/well Expi293F cells were added at 2.5 × 10E6 cells/mL. The culture supernatant was harvested and spun for 10 min at 200 × g to remove cells and cellular debris. The clarified supernatant was subsequently sterile-filtered using a 0.22 μm vacuum filter and stored at 4°C until use. For crystallization, the protein was produced in HEK293 GnTI−/−cells via transient transfection with 10% of Furin-pCDNA2004.

### Purification of 459 V2-SET gp140 trimers using Lectin and SEC

The recombinant 459C V2-SET Envs were purified by a 2-step purification protocol applying a Galantus nivalis-lectin column (Vectorlabs) for the initial purification and subsequently a Superdex200 Increase column (GE) for the polishing step to remove residual contaminants. For the lectin step, the culture supernatant was diluted with 40 mM Tris, 500 mM NaCl pH7.5, and passed over a 4 mL CV Tricorn 10–50 lectin agarose column at 300 cm/hr. Subsequently, the column was washed with 4 column volumes (CV) of 40 mM Tris, 500 mM NaCl pH7.5, and eluted with 4 CV of 40 mM Tris, 500 mM NaCl, 1 M mannopyranoside pH 7.5 with an upflow of 120 cm/hr. The eluate was concentrated using a spin concentrator (50 K, Amicon Ultra, Millipore) and the protein was further purified using a Superdex200 Increase 10/300 column using 20 mM citrate, 75 mM NaCl, 5% sucrose, and 0.03% Tween 80 pH 6.0 as running buffer. The second peak contained the HIV gp140 trimer. The fractions containing this peak were pooled, and the protein concentration was determined using OD280 and stored a 4°C until use.

### Size Exclusion Chromatography and Multi-angle Light Scattering Analysis

Purity and mass were confirmed by size exclusion chromatography (SEC) and multi-angle light scattering (MALS) using a high-performance liquid chromatography system (Agilent Technologies) and miniDAWN TREOS (Wyatt) instrument coupled to an Optilab T-rEX Refractive Index Detector (Wyatt). In total, 40 μg of purified protein was applied to a TSK-Gel G3000SWxl column (Tosoh Bioscience) equilibrated in running buffer (150 mM sodium phosphate, 50 mM sodium chloride, pH 7.0) at 1 mL/min. In other experiments, analysis was performed on supernatants instead of purified Env. The data were analyzed using the Astra 6 software package, and molecular weight calculations were derived from the refractive index signal.

### Bio-Layer Interferometry (Octet)

Antibodies were immobilized on anti-hIgG (AHC) sensors (FortéBio) at a concentration of 1 μg/mL in 10× kinetics buffer (FortéBio) in 96-half well black flat bottom polypylene microplates (Forté-Bio). Experiments were performed on an Octet RED384 instrument (Pall-FortéBio) at 30°C, shaking speed 1,000 rpm. Activation was 300 s, immobilization of antibodies 600 s, washing 300 s, and binding the Envs 1,200 s, followed by a dissociation of 1,200 s, all shaking at 1,000 rpm. The data analysis was performed using the FortéBio Data Analysis 8.1 software (FortéBio).

### 459C trimer probe and H703, CT184, T250.4 DS SOSIP trimer probe production

AVI-tagged H703, CT184, T250.4 repaired and stabilized (RnS) DS SOSIP trimers, and 459C SET trimers, were produced using transient transfection in FreeStyle 293F cells (Thermo Fisher Scientific). In brief, pre-mixed 600 mg constructs encoding N-terminus HRV3C cleavable single-chain Fc-tagged and C-terminus AVI-tagged trimers and 150 mg furin plasmid in 2.25 ml Turbo293 transfection reagent (Speed BioSystems) were added to 0.8 liter of cells at a cell density of 2 × 10^6^ viable cells/ml. Transfected cells were cultured for 6 days in an orbital shaker at 125 rpm at 37°C in a humidified 9% CO_2_ incubator before the supernatant was harvested.

Subsequently, the supernatant was incubated with 5 ml of PBS-equilibrated protein A resin for 1-2 h. The trimer-bound resin was then washed with PBS, collected, and then incubated at 4°C overnight in a 3-ml mixture comprised of 200 μg of HRV3C and BirA biotin-protein ligase mixture (Avidity) according to manufacturer instructions. The HRV3C-liberated and biotinylated trimers were applied to a Superdex 200 16/600 gel filtration column equilibrated with PBS the next day. Peak fractions corresponding to trimers were pooled and negatively selected using a V3 cocktail column containing six V3-directed antibodies (1006-15D, 2219, 2557, 2558, 3074, and 50.1).

### Trimer ELISA for serum samples and monoclonal antibodies

The ELISA using the autologous 459C trimer was performed by coating 96-well Costar half plates (Costar High Binding Half-Area; Corning, Kennebunk, ME) with 2 ug/ml streptavidin overnight at 4°C. The plates were then washed and coated with biotinylated 459C WT, OPT and ALT trimers. For heterologous BG505 DS-SOSIP trimer binding, the method was modified based on previously reported method with lectin captured trimer (Cheng et al., 2020). Briefly, 96-well Costar half plates were coated with 2 mg/ml Lectin and after a one-hour blocking step with 5% skim milk/PBS, the BG505 trimer or glycan-base trimer was incubated on the plates for two hours at RT. After washing the plates 5x with PBS-T, serially diluted sera (at various starting dilutions) or monoclonal antibodies at 2 mg/ml was added to the plates and further incubated for one hour at RT. The plates were then washed 5x with PBS-T and goat anti-rhesus HRP-conjugated secondary antibody was added to the plates at a 1:5000 dilution for one hour at RT. The plates were then developed for 10 minutes using tetramethylbenzidine (TMB) substrate (SureBlue; KPL, Gaithersburg, MD). To stop the reaction, 1 N H_2_SO_4_ sulfuric acid was added. The plates were then read at OD450 nm and values were documented. EC50 for serum binding was calculated when all of the bindings reached the peak OD value, otherwise AUC or endpoint titers were calculated if not all samples reached the peak values.

### Anti-Fusion-Peptide (FP8) ELISA

The ELISA for assessing the response to Fusion-Peptide (FP8) was conducted according to a previously described method (Cheng et al., 2020). Biotinylated eight-residue fusion peptide (FP8-PEG-biotin) was coated in 96-well streptavidin-coated plates (Thermo Fisher) overnight. Subsequently, the plates were blocked with B3T buffer (comprising 150 mM NaCl, 50 mM Tris-HCl, 1 mM EDTA, 3.3% fetal bovine serum, 2% bovine albumin, 0.07% Tween 20, and 0.02% thimerosal). The serum was serially diluted at 7-point 5-fold dilution and incubated for 1hr. Goat anti-rhesus HRP conjugated secondary was added and incubated for an hour. The plates were then developed for 10 minutes using tetramethylbenzidine (TMB) substrate (SureBlue; KPL, Gaithersburg, MD). To stop the reaction, 1 N H_2_SO_4_ sulfuric acid was added. Finally, the plates were read on a microplate spectrophotometer (Biotek Epoch, Winooski, VT) to determine the endpoint titer for each sample.

### Neutralization Assays

Neutralization assays using a single round of infection Env-pseudovirus were performed using TZM-Bl target cells and heat-inactivated sera(Kong et al., 2016). Briefly, 293T cells were cotransfected with an Env expression plasmid and a sPG3ΔEnv backbone to generate the Env-pseudovirus stocks used in neutralization assays. The sera were assessed at various dilutions, using an 8-point 4-fold dilution method which began at a dilution factor of 1:20. The 50% inhibitory dilutions (ID50) were used to assess breadth across the 10 FP-sensitive HIV-1 strains and to test for any correlative relationship with the plasma and B cell parameters tested with Prism.

### HIV-1 envelope trimer immunogens

All HIV-1 envelope trimer immunogens were prepared using transiently transfected 293F cells as previously described (Xu et al., 2018). Broadly neutralizing antibodies 2G12 or VRC01 were used in affinity chromatography to purify the trimers, along with gel filtration (Superdex200 16/60HL column) and a 447-52D affinity column functioning as a negative selection to remove V3-exposed trimers. Antigenicity tests were performed on the trimers with or without mixing with adjuvant using a Meso Scale Discovery (MSD) platform as previously described.

### Isolation of rhesus monoclonal antibodies

Cryopreserved rhesus macaque PBMCs were thawed and stained with LIVE/DEAD fixable violet dead cell stain (Life Technologies). After washing, cells were stained with a cocktail of anti-human antibodies, including CD3 (clone SP34-2; BD Biosciences), CD4 (clone OKT4; BioLegend), CD8 (clone RPA-T8; BioLegend), CD14 (clone M5E2; BioLegend), CD20 (clone 2H7; BioLegend), IgG (G18-145;BD Biosciences), and IgM (clone G20-127; BD Biosciences), and with fluorescently labeled trimer probe (BG505 DS-SOSIP.avi) and peptide probe (FP9-PEG12-biotin) (Kong et al., 2016). Vivid-CD3-CD4-CD8-CD14-CD20+IgG+IgM-memory B cells that are positively stained with both trimer and peptide probes were sorted into 96-well plates containing lysis solution as previously described (Kong et al., 2016). Nested PCR was performed using published primers (Mason et al., 2016; Sundling et al., 2012). Heavy and light chain sequences were cloned into expression vectors containing rhesus macaque immunoglobulin constant regions. IgG was expressed by cotransfecting Expi293FTM cells with equal amounts of paired heavy and light chain plasmids and purified using protein A Fast Flow (GE Healthcare) according to the manufacturer’s instructions.

### Rapid assembly, transfection, and production of immunoglobulins (RATP-Ig)

Part of the antigen-specific sorted B cells were subjected to RATP-Ig analysis as previously described(Lima et al., 2022). Single-cell RNA was purified with RNAclean beads (Beckman Coulter). cDNA was then synthesized using 5’ RACE reverse-transcription and amplified by PCR. An aliquot of enriched cDNA was sequenced using 2×150 paired-end reads on an Illumina MiSeq. For immunoglobulin production, enriched variable regions were assembled into expression cassettes that include CMV, and HC/LC-TBGH polyA fragments. Assembled cassettes were amplified by PCR and transfected into Expi293 cells in 96-well deep-well plates using the Expi293 Transfection Kit (ThermoFisher Scientific). Cell cultures were grown at 37C and 8% CO2, with 1100 RPM shaking for 5–7 days. Cell culture supernatants were harvested by centrifugation.

### Antibody gene assignment, nomenclature and genetic analysis

Antibody sequences were annotated using IMGT V/Quest and IgBlast (Ye et al., 2013) against the donor-specific gene database. Antibodies sharing both heavy and light chain V and J gene assignments and junction region were grouped into one lineage. Monoclonal antibodies were named to include information from the donor and the sorting plate and well containing the single B-cell. For example, the designation mAb J601-1B2, indicates a mAb isolated from well B2 on sorting plate 1 from animal J601. For both heavy chain and kappa chains, germline V genes from a public database (Ramesh et al., 2017) and genes predicted to be used by FP antibodies but specific to an animal were combined to construct a phylogenetic tree using MEGA6 with GTR+G substitution model (Tamura et al., 2013).

### Antibody preparation

The antibody heavy– and light-chain variable region DNA sequences were synthesized and separately cloned into an mammalian cell expression plasmid and sequenced. Plasmids encoding heavy and light chain pairs were co-transfected into Expi293F cells (Thermo Fisher) as described previously. The cells were grown for six days, and the culture supernatants were harvested and loaded on protein A columns. After washing the columns with PBS, the IgG proteins were eluted with a low pH buffer. For antibody Fab preparation, an HRV3C cleavage site was inserted in the heavy-chain hinge region, and the purified IgG protein was digested with HRV3C. The Fab was separated from Fc by passing the digestion mixture through a protein A column and then further purified through a Superdex 200 column (Cytiva).

### Negative-stain EM analysis

The purified antibody Fab was mixed with HIV-1 Env trimer at a 3:1 molar ratio. The mixture was diluted with a negative-stain buffer (10 mM HEPES, pH 7.0, and 150 mM NaCl) to approximately 0.02 mg/ml, and the diluted Fab-Env mixture was transferred to a freshly glow-discharged carbon-coated grid and incubated for a few seconds, and the liquid was removed with a piece of filter paper. The grid was rinsed 3 times with the negative-stain buffer and then stained with 0.75% uranyl formate for 30 seconds. Data were collected using a Talos F200C transmission electron microscope (ThermoFisher Scientific) operated at 200 kV. Particle picking, 2D classification, and 3D reconstruction were carried out using CryoSparc 4.6. The 3D volumes of the Fab-Env complex density were visualized and analyzed, and figures were prepared using UCSF ChimeraX.

### Cryo-EM structure determination

To prepare cryo-EM grids, purified Env trimers at 1-4 mg/ml were mixed with antibody Fab; n-Dodecyl β-D-maltoside (DDM) was added to a final concentration of 0.1mM, and the mixture was applied to a freshly glow discharged carbon-coated copper grid (CF 1.2/1.3 300 mesh) or a Quantifoil-gold 2/2 holey carbon grid. The sample was vitrified in liquid ethane using a Vitrobot Mark IV. Cryo-EM data were collected using Leginon software on a Titan Krios electron microscope operating at 300 kV, equipped with a Gatan K3-BioQuantum direct detection device. Exposures were taken with a total electron fluence of 58.06 e-/Å^2^. The total dose was fractionated for 2.5 s over 50 raw frames. Some cryo-EM data were acquired on a Titan Krios operating at 300 kV, equipped with a K2 Summit detector (Gatan) operating in counting mode and using SerialEM 4.0. The dose was fractionated over 40 raw frames. The movie frames were aligned and dose-weighted, and CTF estimation, particle picking, 2D classifications, *ab initio* model generation, heterogeneous refinements, homogeneous 3D refinements, non-uniform refinement, and local resolution calculations were carried out using cryoSPARC 4.4.

A model of the antibody Fab was generated using AlphaFold2 or AlphaFold3. The Fab model and the structure of BG505 DS-SOSIP (PDB: 6cdi)(Xu et al., 2018) were docked into the cryo-EM map using UCSF ChimeraX. This initial model was then manually rebuilt to fit into the density and match the sequence of the Env trimer of the specific HIV-1 strain using Coot. The structural models were refined using Phenix. Overall structures were evaluated using MolProbity. Protein interface calculations were performed using PISA. Structural figures were generated using PyMOL (Schrodinger; http://www.pymol.org) and UCSF ChimeraX.

## QUANTIFICATION AND STATISTICAL ANALYSES

A two-tailed Mann-Whitney t-test comparing the mean and standard error of the mean (SEM) was performed when comparing two groups in serologic and B-cell studies. For multiple comparison analysis, a Kruskal-Wallis test followed by the two-stage linear step-up procedure of Benjamini, Krieger and Yekutieli tests was performed to assess differences in multiple groups. A two-tailed Pearson correlation coefficient test was used to assess correlations between plasma and B cell parameters and neutralization titers and breadth, when data obeyed normality assumptions. When the normality assumption failed, a 2-tailed Spearman correlation coefficient test was performed. *: p<0.05, **: p<0.01, ***: p<0.0001.

## REFERENCES

1. Bbosa, N., Kaleebu, P., and Ssemwanga, D. (2019). HIV subtype diversity worldwide. Curr Opin HIV AIDS 14, 153–160.

2. Bianchi, M., Turner, H.L., Nogal, B., Cottrell, C.A., Oyen, D., Pauthner, M., Bastidas, R., Nedellec, R., McCoy, L.E., Wilson, I.A., et al. (2018). Electron-Microscopy-Based Epitope Mapping Defines Specificities of Polyclonal Antibodies Elicited during HIV-1 BG505 Envelope Trimer Immunization. Immunity 49, 288–300.e288.

3. Bonsignori, M., Montefiori, D.C., Wu, X., Chen, X., Hwang, K.K., Tsao, C.Y., Kozink, D.M., Parks, R.J., Tomaras, G.D., Crump, J.A., et al. (2012). Two distinct broadly neutralizing antibody specificities of different clonal lineages in a single HIV-1-infected donor: implications for vaccine design. J Virol 86, 4688–4692.

4. Bricault, C.A., Yusim, K., Seaman, M.S., Yoon, H., Theiler, J., Giorgi, E.E., Wagh, K., Theiler, M., Hraber, P., Macke, J.P., et al. (2019). HIV-1 Neutralizing Antibody Signatures and Application to Epitope-Targeted Vaccine Design. Cell Host Microbe 25, 59–72.e58.

5. Cale, E.M., Driscoll, J.I., Lee, M., Gorman, J., Zhou, T., Lu, M., Geng, H., Lai, Y.T., Chuang, G.Y., Doria-Rose, N.A., et al. (2022). Antigenic analysis of the HIV-1 envelope trimer implies small differences between structural states 1 and 2. J Biol Chem 298, 101819.

6. Cale, E.M., Gorman, J., Radakovich, N.A., Crooks, E.T., Osawa, K., Tong, T., Li, J., Nagarajan, R., Ozorowski, G., Ambrozak, D.R., et al. (2017). Virus-like Particles Identify an HIV V1V2 Apex-Binding Neutralizing Antibody that Lacks a Protruding Loop. Immunity 46, 777–791.e710.

7. Cale, E.M., Shen, C.H., Olia, A.S., Radakovich, N.A., Rawi, R., Yang, Y., Ambrozak, D.R., Bennici, A.K., Chuang, G.Y., Crooks, E.D., et al. (2024). A multidonor class of highly glycan-dependent HIV-1 gp120-gp41 interface-targeting broadly neutralizing antibodies. Cell Rep 43, 115010.

8. Cheng, C., Duan, H., Xu, K., Chuang, G.Y., Corrigan, A.R., Geng, H., O’Dell, S., Ou, L., Chambers, M., Changela, A., et al. (2020). Immune Monitoring Reveals Fusion Peptide Priming to Imprint Cross-Clade HIV-Neutralizing Responses with a Characteristic Early B Cell Signature. Cell Rep 32, 107981.

9. Cheng, C., Xu, K., Kong, R., Chuang, G.Y., Corrigan, A.R., Geng, H., Hill, K.R., Jafari, A.J., O’Dell, S., Ou, L., et al. (2019). Consistent elicitation of cross-clade HIV-neutralizing responses achieved in guinea pigs after fusion peptide priming by repetitive envelope trimer boosting. PLoS One 14, e0215163.

10. Corrigan, A.R., Duan, H., Cheng, C., Gonelli, C.A., Ou, L., Xu, K., DeMouth, M.E., Geng, H., Narpala, S., O’Connell, S., et al. (2021). Fusion peptide priming reduces immune responses to HIV-1 envelope trimer base. Cell Rep 35, 108937.

11. Cottrell, C.A., van Schooten, J., Bowman, C.A., Yuan, M., Oyen, D., Shin, M., Morpurgo, R., van der Woude, P., van Breemen, M., Torres, J.L., et al. (2020). Mapping the immunogenic landscape of near-native HIV-1 envelope trimers in non-human primates. PLoS Pathog 16, e1008753.

12. Davenport, T.M., Friend, D., Ellingson, K., Xu, H., Caldwell, Z., Sellhorn, G., Kraft, Z., Strong, R.K., and Stamatatos, L. (2011). Binding interactions between soluble HIV envelope glycoproteins and quaternary-structure-specific monoclonal antibodies PG9 and PG16. J Virol 85, 7095–7107.

13. de Taeye, S.W., Ozorowski, G., Torrents de la Peña, A., Guttman, M., Julien, J.P., van den Kerkhof, T.L., Burger, J.A., Pritchard, L.K., Pugach, P., Yasmeen, A., et al. (2015). Immunogenicity of Stabilized HIV-1 Envelope Trimers with Reduced Exposure of Non-neutralizing Epitopes. Cell 163, 1702–1715.

14. Duan, H., Corrigan, A.R., Cheng, C., Biju, A., Gonelli, C.A., Olia, A.S., Teng, I.T., Xu, K., O’Dell, S., Narpala, S., et al. (2024). Long trimer-immunization interval and appropriate adjuvant reduce immune responses to the soluble HIV-1-envelope trimer base. iScience 27, 108877.

15. Ferguson, M.R., Rojo, D.R., von Lindern, J.J., and O’Brien, W.A. (2002). HIV-1 replication cycle. Clin Lab Med 22, 611–635.

16. Gorman, J., Chuang, G.Y., Lai, Y.T., Shen, C.H., Boyington, J.C., Druz, A., Geng, H., Louder, M.K., McKee, K., Rawi, R., et al. (2020). Structure of Super-Potent Antibody CAP256-VRC26.25 in Complex with HIV-1 Envelope Reveals a Combined Mode of Trimer-Apex Recognition. Cell Rep 31, 107488.

17. Gristick, H.B., Hartweger, H., Nishimura, Y., Gavor, E., Nagashima, K., Koranda, N.S., Gnanapragasam, P.N.P., Kakutani, L.M., Segovia, L., Donau, O., et al. (2025). Design and characterization of HIV-1 vaccine candidates to elicit antibodies targeting multiple epitopes. bioRxiv.

18. Hodge, E.A., Naika, G.S., Kephart, S.M., Nguyen, A., Zhu, R., Benhaim, M.A., Guo, W., Moore, J.P., Hu, S.L., Sanders, R.W., et al. (2022). Structural dynamics reveal isolate-specific differences at neutralization epitopes on HIV Env. iScience 25, 104449.

19. Hu, J.K., Crampton, J.C., Cupo, A., Ketas, T., van Gils, M.J., Sliepen, K., de Taeye, S.W., Sok, D., Ozorowski, G., Deresa, I., et al. (2015). Murine Antibody Responses to Cleaved Soluble HIV-1 Envelope Trimers Are Highly Restricted in Specificity. J Virol 89, 10383–10398.

20. Huang, J., Kang, B.H., Pancera, M., Lee, J.H., Tong, T., Feng, Y., Imamichi, H., Georgiev, I.S., Chuang, G.Y., Druz, A., et al. (2014). Broad and potent HIV-1 neutralization by a human antibody that binds the gp41-gp120 interface. Nature 515, 138–142.

21. Huang, J., Ofek, G., Laub, L., Louder, M.K., Doria-Rose, N.A., Longo, N.S., Imamichi, H., Bailer, R.T., Chakrabarti, B., Sharma, S.K., et al. (2012). Broad and potent neutralization of HIV-1 by a gp41-specific human antibody. Nature 491, 406–412.

22. Kong, R., Duan, H., Sheng, Z., Xu, K., Acharya, P., Chen, X., Cheng, C., Dingens, A.S., Gorman, J., Sastry, M., et al. (2019). Antibody Lineages with Vaccine-Induced Antigen-Binding Hotspots Develop Broad HIV Neutralization. Cell 178, 567–584.e519.

23. Kong, R., Xu, K., Zhou, T., Acharya, P., Lemmin, T., Liu, K., Ozorowski, G., Soto, C., Taft, J.D., Bailer, R.T., et al. (2016). Fusion peptide of HIV-1 as a site of vulnerability to neutralizing antibody. Science 352, 828–833.

24. Kulp, D.W., Steichen, J.M., Pauthner, M., Hu, X., Schiffner, T., Liguori, A., Cottrell, C.A., Havenar-Daughton, C., Ozorowski, G., Georgeson, E., et al. (2017). Structure-based design of native-like HIV-1 envelope trimers to silence non-neutralizing epitopes and eliminate CD4 binding. Nat Commun 8, 1655.

25. Lima, N.S., Musayev, M., Johnston, T.S., Wagner, D.A., Henry, A.R., Wang, L., Yang, E.S., Zhang, Y., Birungi, K., Black, W.P., et al. (2022). Primary exposure to SARS-CoV-2 variants elicits convergent epitope specificities, immunoglobulin V gene usage and public B cell clones. Nat Commun 13, 7733.

26. Martinez-Murillo, P., Tran, K., Guenaga, J., Lindgren, G., Àdori, M., Feng, Y., Phad, G.E., Vázquez Bernat, N., Bale, S., Ingale, J., et al. (2017). Particulate Array of Well-Ordered HIV Clade C Env Trimers Elicits Neutralizing Antibodies that Display a Unique V2 Cap Approach. Immunity 46, 804–817.e807.

27. Mogus, A.T., Liu, L., Jia, M., Ajayi, D.T., Xu, K., Kong, R., Huang, J., Yu, J., Kwong, P.D., Mascola, J.R., et al. (2020). Virus-Like Particle Based Vaccines Elicit Neutralizing Antibodies against the HIV-1 Fusion Peptide. Vaccines (Basel) 8.

28. Munro, J.B., and Lee, K.K. (2018). Probing Structural Variation and Dynamics in the HIV-1 Env Fusion Glycoprotein. Curr HIV Res 16, 5–12.

29. Olia, A.S., Cheng, C., Zhou, T., Biju, A., Harris, D.R., Changela, A., Duan, H., Ivleva, V.B., Kong, W.P., Ou, L., et al. (2023). Soluble prefusion-closed HIV-envelope trimers with glycan-covered bases. iScience 26, 107403.

30. Ou, L., Kong, W.P., Chuang, G.Y., Ghosh, M., Gulla, K., O’Dell, S., Varriale, J., Barefoot, N., Changela, A., Chao, C.W., et al. (2020). Preclinical Development of a Fusion Peptide Conjugate as an HIV Vaccine Immunogen. Sci Rep 10, 3032.

31. Pratap, P.P., Cottrell, C.A., Quinn, J., Carnathan, D.G., Bader, D.L.V., Tran, A.S., Enemuo, C.A., Ngo, J.T., Richey, S.T., Gao, H., et al. (2025). Immunofocusing on the conserved fusion peptide of HIV envelope glycoprotein in rhesus macaques. bioRxiv.

32. Read, B.J., Won, L., Kraft, J.C., Sappington, I., Aung, A., Wu, S., Bals, J., Chen, C., Lee, K.K., Lingwood, D., et al. (2022). Mannose-binding lectin and complement mediate follicular localization and enhanced immunogenicity of diverse protein nanoparticle immunogens. Cell Rep 38, 110217.

33. Roark, R.S., Habib, R., Gorman, J., Li, H., Connell, A.J., Bonsignori, M., Guo, Y., Hogarty, M.P., Olia, A.S., Sowers, K.J., et al. (2025). Structural and genetic basis of HIV-1 envelope V2 apex recognition by rhesus broadly neutralizing antibodies. J Exp Med 222.

34. Rutten, L., Lai, Y.T., Blokland, S., Truan, D., Bisschop, I.J.M., Strokappe, N.M., Koornneef, A., van Manen, D., Chuang, G.Y., Farney, S.K., et al. (2018). A Universal Approach to Optimize the Folding and Stability of Prefusion-Closed HIV-1 Envelope Trimers. Cell Rep 23, 584–595.

35. Sanders, R.W., Derking, R., Cupo, A., Julien, J.P., Yasmeen, A., de Val, N., Kim, H.J., Blattner, C., de la Peña, A.T., Korzun, J., et al. (2013). A next-generation cleaved, soluble HIV-1 Env trimer, BG505 SOSIP.664 gp140, expresses multiple epitopes for broadly neutralizing but not non-neutralizing antibodies. PLoS Pathog 9, e1003618.

36. Setliff, I., Shiakolas, A.R., Pilewski, K.A., Murji, A.A., Mapengo, R.E., Janowska, K., Richardson, S., Oosthuysen, C., Raju, N., Ronsard, L., et al. (2019). High-Throughput Mapping of B Cell Receptor Sequences to Antigen Specificity. Cell 179, 1636–1646.e1615.

37. Stewart-Jones, G.B., Soto, C., Lemmin, T., Chuang, G.Y., Druz, A., Kong, R., Thomas, P.V., Wagh, K., Zhou, T., Behrens, A.J., et al. (2016). Trimeric HIV-1-Env Structures Define Glycan Shields from Clades A, B, and G. Cell 165, 813–826.

38. Voss, J.E., Andrabi, R., McCoy, L.E., de Val, N., Fuller, R.P., Messmer, T., Su, C.Y., Sok, D., Khan, S.N., Garces, F., et al. (2017). Elicitation of Neutralizing Antibodies Targeting the V2 Apex of the HIV Envelope Trimer in a Wild-Type Animal Model. Cell Rep 21, 222–235.

39. Wang, X., Cottrell, C.A., Hu, X., Ray, R., Bottermann, M., Villavicencio, P.M., Yan, Y., Xie, Z., Warner, J.E., Ellis-Pugh, J.R., et al. (2024). mRNA-LNP prime boost evolves precursors toward VRC01-like broadly neutralizing antibodies in preclinical humanized mouse models. Sci Immunol 9, eadn0622.

40. Willis, J.R., Berndsen, Z.T., Ma, K.M., Steichen, J.M., Schiffner, T., Landais, E., Liguori, A., Kalyuzhniy, O., Allen, J.D., Baboo, S., et al. (2022). Human immunoglobulin repertoire analysis guides design of vaccine priming immunogens targeting HIV V2-apex broadly neutralizing antibody precursors. Immunity 55, 2149–2167.e2149.

41. Xu, K., Acharya, P., Kong, R., Cheng, C., Chuang, G.Y., Liu, K., Louder, M.K., O’Dell, S., Rawi, R., Sastry, M., et al. (2018). Epitope-based vaccine design yields fusion peptide-directed antibodies that neutralize diverse strains of HIV-1. Nat Med 24, 857–867.

