## Supplementary data for "Vaccine Elicitation of HIV-1 Neutralizing Antibodies Against Both V2 Apex and Fusion Peptide in Rhesus Macaques"

**A**

| | 33 | 49 | 130 | 132 | 133 | 156 | 158 | 160 | 161 | 162 | 163 | 164 | 165 | 166 | 167 | 168 | 169 | 170 | 171 | 172 | 173 | 177 | 223 | 297 | 306 | 322 | 335 | 636 | 644 | 677 |
|---|---|---|---|---|---|---|---|---|---|---|---|---|---|---|---|---|---|---|---|---|---|---|---|---|---|---|---|---|---|---|
| 459C WT | N | K | N | T | N | N | S | N | A | T | T | E | I | R | D | R | K | K | E | M | Y | Y | F | T | S | D | E | N | R | N |
| 459C OPT | N | K | E | T | A | N | S | N | M | T | T | E | L | R | D | K | K | K | K | V | S | Y | Y | I | S | D | K | S | R | N |
| 459C ALT | N | E | D | K | A | N | T | N | I | T | T | S | V | K | G | K | R | Q | Q | E | H | Y | Y | V | S | E | K | S | R | N |

**B**

| Rns modification | | 155 | 201 | 302 | 304 | 318 | 320 | 514 | 535 | 538 | 567 | 568 | 579 | 632 | 636 | 639 |
|---|---|---|---|---|---|---|---|---|---|---|---|---|---|---|---|---|
| | SOSIP | Q | L | N | V | Y | N | I | L | I | K | D | A | N | K | N |
| | Rns | K | I | M | R | A | L | N | P | P | E | V | G | F | I | V |

**C** RnS 459C production efficiency

| 459C | Yield (mg/L) | Purification method |
|---|---|---|
| WT | 25.6 | Lectin/SEC/14E |
| OPT | 35.3 | VRC26/SEC/14E |
| ALT | 25.8 | Lectin/SEC/14E |

**D**

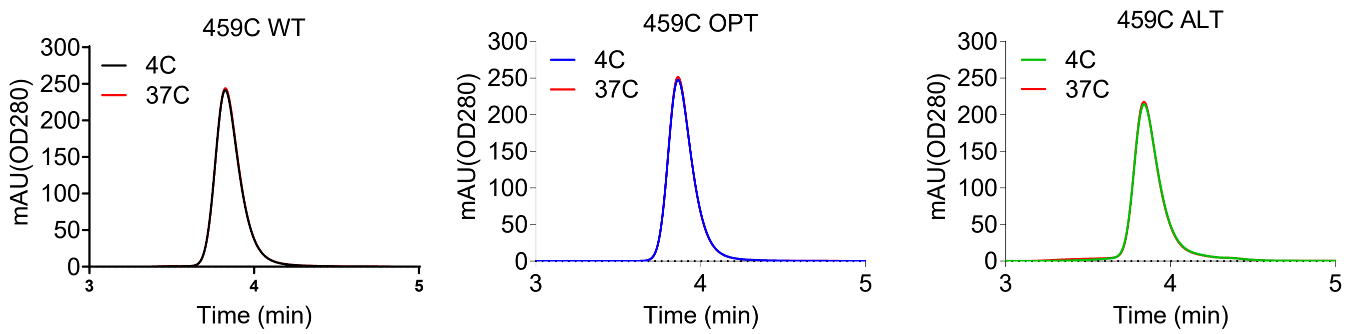

**E**

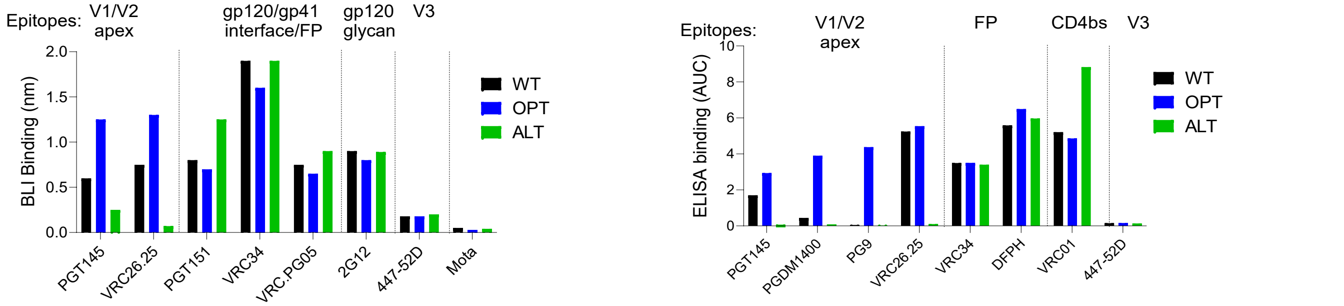

**F**

RMSD of pairwise aligned Cα atoms (Å)

| | WT | ALT | OPT |
|---|---|---|---|
| WT | | 1.60 | 1.06 |
| ALT | 1586 | | 2.21 |
| OPT | 1544 | 1638 | |

Number of the aligned Cα atoms in Pymol

**G**

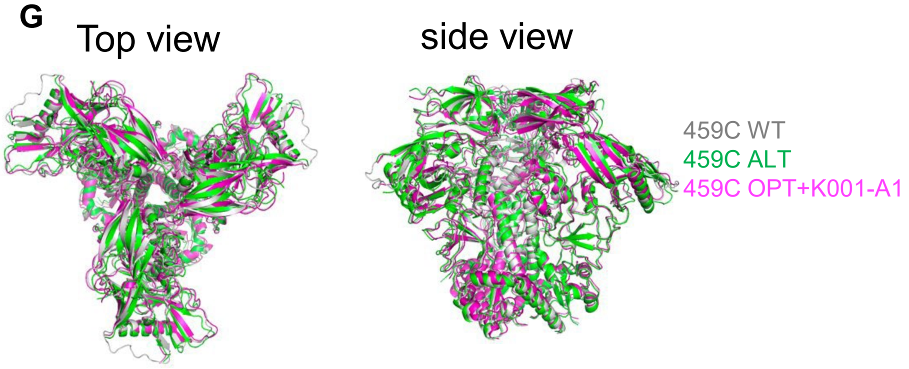

**Figure S1. RnS V2-SET modification confers stable prefusion conformation and higher V2-specific bNab binding, related to Figure 1.** **(A, B)** 459C WT, OPT and ALT trimers were made with V2-SET (**A**) and Rns modifications (**B**). (**C**) 459C WT, OPT and ALT trimers produced well. **(D)** 459C V2-SET trimers stay prefusion conformation at 37 degree for 4 weeks. **(E)** Binding of 459C WT, OPT and ALT trimers to HIV-specific mAbs in BLI (left) or ELISA (right). (**F**) RMSD of pairwise aligned Ca atoms between 459C WT, OPT and ALT trimers. **(G)** Structure alignment of 459C WT, OPT and ALT trimers.

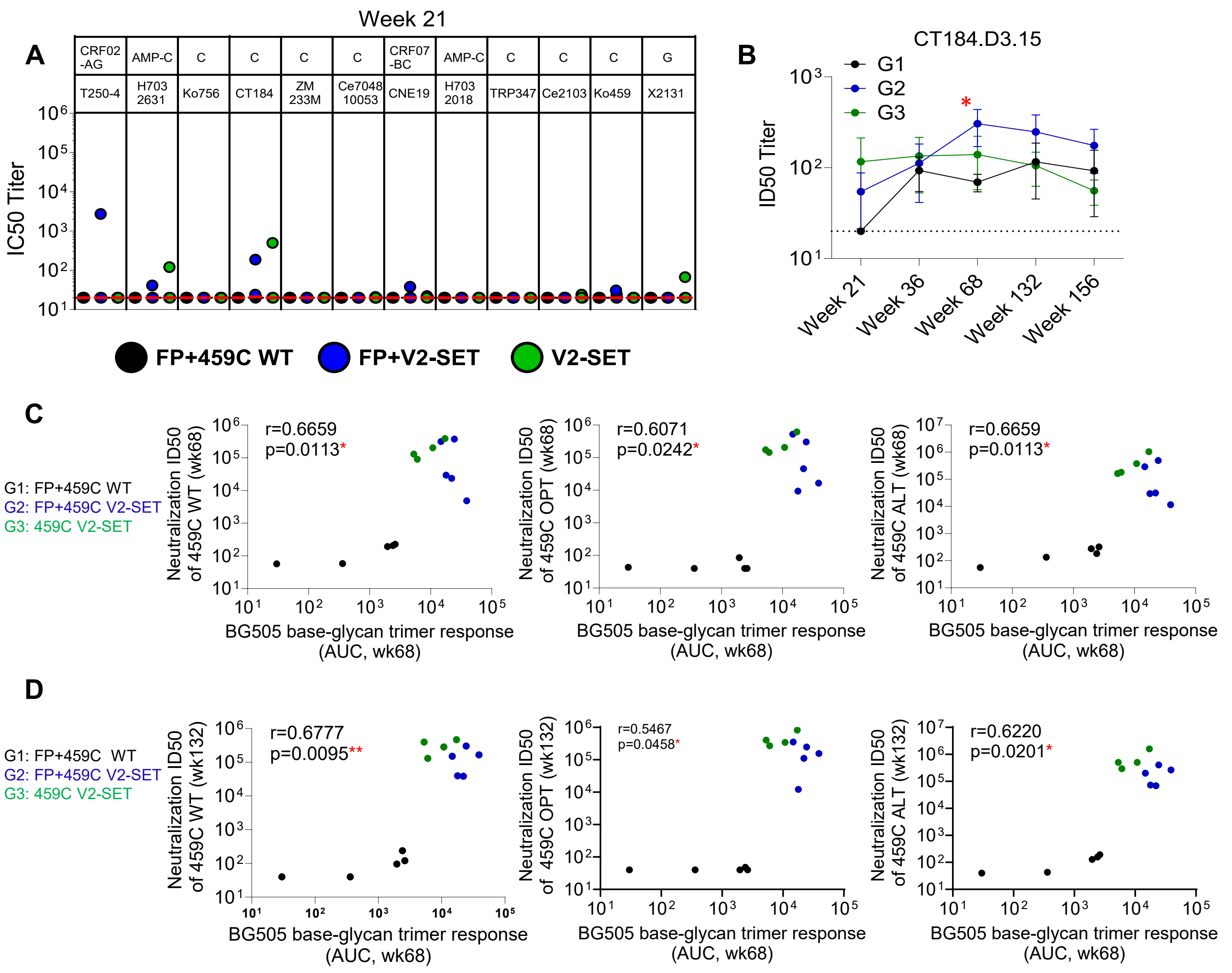

**Figure S2. A. V2-SET modification confer earlier and higher heterologous neutralizing activity and higher plasma non-base trimer responses, related to Figure 2 and 3. (A)** Serum neutralization ID50 titers at wk21 on tier 2 heterologous viruses. (**B**) Serum ID50 titers on tier 2 clade C CT184.D3.15 virus at wks 21, 36, 68, 132 and 156 after immunization. (**C,D**) Plasma responses to glycan-base BG505 DS-SOSIP trimer at wk68 correlate with ID50 titers on 459C WT, OPT and ALT viruses at wk68 (**C**) and wk132 (**D**). Animals from G1, G2, G3 are in black, blue and green fonts/dots, respectively. Spearman 2-tailed correlation coefficient test were used for the correlation analysis. *: p<0.05; **: p <0.01.

| | | | | Wk21 Serum ID50 Titer in TZM.bl cells (1/x) | | | | | | | | | | |
|---|---|---|---|---|---|---|---|---|---|---|---|---|---|---|
| Animal # | Group | Timepoint | T250-4 | H703_2631 | Ko756_38_Tb12 | CT184_D3.15 | ZM233M.PB6 | Ce704810053_2B7 | CNE19 | H703_2018 | TRP347.2.00_B1_1 | Ce2103_E8 | Ko459_T68_4 | X2131_C1_B5 |
| G66C | FP+459C | Week 21 | <20 | <20 | <20 | <20 | <20 | <20 | <20 | <20 | <20 | <20 | <20 | <20 |
| G66V | FP+459C | Week 21 | <20 | <20 | <20 | <20 | <20 | <20 | <20 | <20 | <20 | <20 | <20 | <20 |
| G67E | FP+459C | Week 21 | <20 | <20 | <20 | <20 | <20 | <20 | <20 | <20 | <20 | <20 | <20 | <20 |
| G70W | FP+459C | Week 21 | <20 | <20 | <20 | <20 | <20 | <20 | <20 | <20 | <20 | <20 | <20 | <20 |
| G71K | FP+459C | Week 21 | <20 | <20 | <20 | <20 | <20 | <20 | <20 | <20 | <20 | <20 | <20 | <20 |
| J399 | FP+SET | Week 21 | <20 | <20 | <20 | <20 | <20 | <20 | <20 | <20 | <20 | <20 | <20 | <20 |
| J400 | FP+SET | Week 21 | <20 | <20 | <20 | <20 | <20 | <20 | <20 | <20 | <20 | <20 | <20 | <20 |
| J601 | FP+SET | Week 21 | 2,723 | 41 | <20 | 21 | <20 | <20 | <20 | <20 | <20 | <20 | 24 | <20 |
| J606 | FP+SET | Week 21 | <20 | <20 | <20 | 187 | <20 | <20 | <20 | <20 | <20 | <20 | 31 | <20 |
| J618 | FP+SET | Week 21 | <20 | <20 | <20 | 24 | <20 | <20 | <20 | <20 | <20 | <20 | <20 | <20 |
| 667 | SET | Week 21 | <20 | <20 | <20 | <20 | <20 | <20 | <20 | <20 | <20 | <20 | <20 | <20 |
| K001 | SET | Week 21 | <20 | 121 | <20 | 501 | <20 | <20 | <20 | <20 | <20 | <20 | <20 | 67 |
| K120 | SET | Week 21 | <20 | <20 | <20 | <20 | <20 | <20 | <20 | <20 | <20 | <20 | <20 | <20 |
| K176 | SET | Week 21 | <20 | <20 | <20 | <20 | <20 | <20 | <20 | <20 | <20 | <20 | <20 | <20 |
| K301 | SET | Week 21 | <20 | <20 | <20 | <20 | <20 | <20 | <20 | <20 | <20 | <20 | <20 | <20 |

| | | | | Wk36 serum ID50 Titer in TZM.bl cells (1/x) | | | | | | | | | | |
|---|---|---|---|---|---|---|---|---|---|---|---|---|---|---|
| Animal # | Group | Timepoint | T250-4 | H703_2631 | Ko756_38_Tb12 | CT184_D3.15 | ZM233M.PB6 | Ce704810053_2B7 | CNE19 | H703_2018 | TRP347.2.00_B1_1 | Ce2103_E8 | Ko459_T68_4 | X2131_C1_B5 |
| G66C | FP+459C | Week 36 | 30 | 45 | <20 | 78 | <20 | <20 | 24 | <20 | <20 | <20 | 43 | <20 |
| G66V | FP+459C | Week 36 | <20 | <20 | <20 | <20 | <20 | <20 | <20 | <20 | <20 | <20 | <20 | <20 |
| G67E | FP+459C | Week 36 | <20 | 43 | <20 | 99 | <20 | <20 | <20 | <20 | <20 | <20 | <20 | <20 |
| G70W | FP+459C | Week 36 | <20 | 43 | <20 | 33 | <20 | <20 | <20 | <20 | <20 | <20 | 23 | <20 |
| G71K | FP+459C | Week 36 | <20 | <20 | <20 | 235 | <20 | <20 | <20 | <20 | <20 | <20 | <20 | <20 |
| J399 | FP+SET | Week 36 | <20 | <20 | <20 | 51 | <20 | <20 | <20 | <20 | <20 | <20 | <20 | <20 |
| J400 | FP+SET | Week 36 | <20 | 60 | <20 | 47 | 33 | 20 | <20 | <20 | <20 | <20 | 35 | <20 |
| J601 | FP+SET | Week 36 | 4,536 | 257 | <20 | 30 | 33 | <20 | <20 | <20 | <20 | <20 | 46 | <20 |
| J606 | FP+SET | Week 36 | <20 | 121 | <20 | 394 | <20 | <20 | <20 | 86 | 23 | <20 | 94 | 70 |
| J618 | FP+SET | Week 36 | 982 | 34 | <20 | 38 | <20 | <20 | <20 | <20 | <20 | <20 | <20 | <20 |
| 667 | SET | Week 36 | 2,440 | 28 | <20 | 30 | <20 | <20 | <20 | <20 | <20 | <20 | <20 | <20 |
| K001 | SET | Week 36 | 1,086 | 119 | <20 | 450 | <20 | <20 | <20 | <20 | 57 | <20 | 53 | 57 |
| K120 | SET | Week 36 | 646 | <20 | <20 | <20 | <20 | <20 | <20 | <20 | <20 | <20 | <20 | <20 |
| K176 | SET | Week 36 | <20 | 47 | <20 | 137 | <20 | <20 | <20 | <20 | 38 | <20 | 26 | <20 |
| K301 | SET | Week 36 | <20 | 102 | <20 | 35 | <20 | <20 | <20 | <20 | <20 | <20 | <20 | 29 |

| | | | | Wk68 serum ID50 Titer in TZM.bl cells (1/x) | | | | | | | | | | |
|---|---|---|---|---|---|---|---|---|---|---|---|---|---|---|
| Animal # | Group | Timepoint | T250-4 | H703_2631 | Ko756_38_Tb12 | CT184_D3.15 | ZM233M.PB6 | Ce704810053_2B7 | CNE19 | H703_2018 | TRP347.2.00_B1_1 | Ce2103_E8 | Ko459_T68_4 | X2131_C1_B5 |
| G66C | FP+459C | Week 68 | <20 | 33 | <20 | 48 | <20 | <20 | 25 | <20 | <20 | <20 | <20 | <20 |
| G66V | FP+459C | Week 68 | <20 | <20 | <20 | <20 | <20 | <20 | <20 | <20 | <20 | <20 | <20 | <20 |
| G67E | FP+459C | Week 68 | <20 | 60 | <20 | 95 | <20 | <20 | 45 | <20 | <20 | <20 | <20 | <20 |
| G70W | FP+459C | Week 68 | <20 | 77 | <20 | 91 | <20 | 43 | 67 | <20 | <20 | <20 | 40 | 45 |
| G71K | FP+459C | Week 68 | <20 | <20 | <20 | 93 | <20 | <20 | <20 | <20 | <20 | <20 | <20 | <20 |
| J399 | FP+SET | Week 68 | 628 | 140 | <20 | 244 | 84 | <20 | <20 | <20 | <20 | <20 | 49 | <20 |
| J400 | FP+SET | Week 68 | 27 | 526 | <20 | 637 | 54 | 31 | 22 | <20 | 38 | <20 | 148 | 38 |
| J601 | FP+SET | Week 68 | 21,249 | 1,026 | 54 | <20 | 89 | <20 | 80 | <20 | <20 | <20 | 67 | <20 |
| J606 | FP+SET | Week 68 | 133 | 125 | <20 | 594 | <20 | <20 | <20 | 61 | <20 | <20 | 98 | 80 |
| J618 | FP+SET | Week 68 | 16,339 | 70 | <20 | 27 | 37 | <20 | <20 | <20 | <20 | <20 | <20 | 22 |
| 667 | SET | Week 68 | 24,511 | 252 | <20 | <20 | <20 | <20 | <20 | <20 | <20 | <20 | 58 | <20 |
| K001 | SET | Week 68 | 11,914 | 159 | <20 | 461 | <20 | <20 | <20 | <20 | 131 | <20 | 79 | 74 |
| K120 | SET | Week 68 | 3,600 | 48 | <20 | <20 | <20 | <20 | <20 | <20 | <20 | <20 | 23 | <20 |
| K176 | SET | Week 68 | <20 | 245 | <20 | 104 | <20 | <20 | <20 | <20 | <20 | <20 | 29 | 28 |
| K301 | SET | Week 68 | 409 | 305 | <20 | 92 | <20 | <20 | <20 | <20 | <20 | <20 | 66 | 22 |

| | | | | Wk132 serum ID50 Titer in TZM.bl cells (1/x) | | | | | | | | | | |
|---|---|---|---|---|---|---|---|---|---|---|---|---|---|---|
| Animal # | Group | Timepoint | T250-4 | H703_2631 | Ko756_38_Tb12 | CT184_D3.15 | ZM233M.PB6 | Ce704810053_2B7 | CNE19 | H703_2018 | TRP347.2.00_B1_1 | Ce2103_E8 | Ko459_T68_4 | X2131_C1_B5 |
| G66C | FP+459C | Week 132 | <20 | 43 | <20 | 59 | <20 | <20 | 58 | <20 | <20 | <20 | 34 | <20 |
| G66V | FP+459C | Week 132 | <20 | 66 | 29 | <20 | <20 | 32 | 90 | <20 | <20 | <20 | <20 | <20 |
| G67E | FP+459C | Week 132 | <20 | 105 | <20 | <20 | <20 | <20 | 100 | <20 | <20 | <20 | <20 | <20 |
| G70W | FP+459C | Week 132 | <20 | 162 | 282 | 393 | <20 | 184 | 259 | 86 | 31 | <20 | 70 | 138 |
| G71K | FP+459C | Week 132 | <20 | <20 | <20 | 87 | <20 | <20 | <20 | <20 | <20 | <20 | <20 | <20 |
| J399 | FP+SET | Week 132 | 421 | 115 | <20 | 724 | 231 | 22 | <20 | <20 | <20 | <20 | <20 | <20 |
| J400 | FP+SET | Week 132 | 710 | 308 | 77 | 354 | 68 | 30 | 58 | <20 | <20 | <20 | 50 | <20 |
| J601 | FP+SET | Week 132 | 13,714 | 797 | 36 | <20 | 165 | 42 | 48 | <20 | <20 | <20 | 43 | <20 |
| J606 | FP+SET | Week 132 | 130 | 25 | <20 | 55 | <20 | <20 | <20 | <20 | <20 | <20 | <20 | <20 |
| J618 | FP+SET | Week 132 | 12,585 | 75 | 61 | 85 | 115 | 52 | 36 | 36 | <20 | 45 | 26 | 23 |
| 667 | SET | Week 132 | NT | NT | NT | NT | NT | NT | NT | NT | NT | NT | NT | NT |
| K001 | SET | Week 132 | 31,680 | 85 | 46 | 190 | <20 | <20 | 44 | 25 | 179 | <20 | 102 | 26 |
| K120 | SET | Week 132 | 4,056 | 38 | <20 | <20 | <20 | <20 | <20 | <20 | <20 | <20 | 39 | <20 |
| K176 | SET | Week 132 | <20 | 341 | 21 | 168 | <20 | <20 | 34 | <20 | 35 | 21 | 37 | 27 |
| K301 | SET | Week 132 | 1,226 | 205 | <20 | 44 | <20 | <20 | <20 | <20 | <20 | <20 | 144 | 25 |

| | | | | Wk156 serum ID50 Titer in TZM.bl cells (1/x) | | | | | | | | | | |
|---|---|---|---|---|---|---|---|---|---|---|---|---|---|---|
| Animal # | Group | Timepoint | T250-4 | H703_2631 | Ko756_38_Tb12 | CT184_D3.15 | ZM233M.PB6 | Ce704810053_2B7 | CNE19 | H703_2018 | TRP347.2.00_B1_1 | Ce2103_E8 | Ko459_T68_4 | X2131_C1_B5 |
| G66C | FP+459C | Week 156 | <20 | 25 | <20 | <20 | <20 | <20 | <20 | <20 | <20 | <20 | <20 | <20 |
| G66V | FP+459C | Week 156 | <20 | 27 | <20 | <20 | <20 | <20 | <20 | <20 | <20 | <20 | <20 | <20 |
| G67E | FP+459C | Week 156 | <20 | <20 | <20 | <20 | <20 | <20 | <20 | <20 | <20 | <20 | <20 | <20 |
| G70W | FP+459C | Week 156 | <20 | 78 | 43 | 345 | <20 | 129 | 106 | 39 | <20 | <20 | 43 | 91 |
| G71K | FP+459C | Week 156 | <20 | <20 | <20 | 57 | <20 | <20 | <20 | <20 | <20 | <20 | <20 | <20 |
| J399 | FP+SET | Week 156 | 45 | 25 | <20 | 495 | 72 | <20 | <20 | <20 | <20 | <20 | <20 | <20 |
| J400 | FP+SET | Week 156 | 187 | 176 | 63 | 241 | 44 | <20 | 21 | <20 | <20 | <20 | 33 | 79 |
| J601 | FP+SET | Week 156 | 18,526 | 2,217 | 64 | <20 | 296 | <20 | 71 | <20 | <20 | <20 | 41 | <20 |
| J606 | FP+SET | Week 156 | 201 | 75 | <20 | 76 | <20 | <20 | <20 | <20 | <20 | <20 | 43 | 62 |
| J618 | FP+SET | Week 156 | 14,559 | 60 | 28 | 46 | 85 | <20 | <20 | <20 | <20 | 37 | <20 | <20 |
| 667 | SET | Week 156 | NT | NT | NT | NT | NT | NT | NT | NT | NT | NT | NT | NT |
| K001 | SET | Week 156 | 16,845 | 36 | <20 | 97 | <20 | <20 | <20 | <20 | 135 | <20 | <20 | <20 |
| K120 | SET | Week 156 | 2,822 | <20 | <20 | <20 | <20 | <20 | <20 | <20 | <20 | <20 | 42 | <20 |
| K176 | SET | Week 156 | <20 | 131 | <20 | 70 | <20 | <20 | 54 | <20 | 33 | <20 | <20 | 38 |
| K301 | SET | Week 156 | 469 | 81 | <20 | 36 | 40 | <20 | <20 | <20 | <20 | <20 | 110 | <20 |

| Color coding for serum neutralization ID50s | | | | | |
|---|---|---|---|---|---|
| | >100,000 | 10,000-100,000 | 1000-10,000 | 100-1000 | 10-100 |

**Figure S3. A. Serum neutralization ID50 titers on heterologous tier 2 viruses at wks 21, 36, 68, 136 and 156, related to Figure 3.** NHPs from G1, G2 and G3 are shown in black, blue and green fonts, respectively in animal # column. ID50 titers are highlighted based on the color coding shown in the bottom of the table.

**A**

IC50 µg/ml

| Antibody | T250-4.SG3 | BG505.W6M.C2.N611Q.SG3 | SIVmac251.30.SG3 |
|---|---|---|---|
| **clade** | **AG** | **A** | |
| J618-P2E7 | >50 | 0.274 | >50 |
| J618-P2H6 | >50 | 20.40 | >50 |
| J618-P2A12 | >50 | 0.162 | >50 |
| J618-P1G4 | >50 | 4.93 | >50 |
| J601-A6 | >50 | 0.351 | >50 |
| K001-B9 | >50 | >50 | >50 |
| K001-B8 | >50 | >50 | >50 |
| VRC34.01 | 0.256 | 0.073 | >50 |

IC80 µg/ml

| Antibody | T250-4.SG3 | BG505.W6M.C2.N611Q.SG3 | SIVmac251.30.SG3 |
|---|---|---|---|
| **clade** | **AG** | **A** | |
| J618-P2E7 | >50 | 1.10 | >50 |
| J618-P2H6 | >50 | >50 | >50 |
| J618-P2A12 | >50 | 22.00 | >50 |
| J618-P1G4 | >50 | >50 | >50 |
| J601-A6 | >50 | 0.525 | >50 |
| K001-B9 | >50 | >50 | >50 |
| K001-B8 | >50 | >50 | >50 |
| VRC34.01 | 1.76 | 0.365 | >50 |

Color coding for antibody neutralization: (µg/ml)

| | |
|---|---|
| <0.001 | .100-1.00 |
| .001-.01 | 1.00-10.0 |
| .01-.100 | 10.0-50.0 |

**B**

| | G70W-2E9 | J601-1B2 | VRC34.01 |
|---|---|---|---|
| **# Viruses** | 208 | 208 | 208 |
| **IC50 <50ug/ml** | 27 | 97 | 105 |
| **IC50 <10ug/ml** | 14 | 82 | 97 |
| **IC50 <1.0ug/ml** | 3 | 13 | 73 |
| **IC50 <0.1ug/ml** | 0 | 1 | 25 |
| **IC50 <0.01ug/ml** | 0 | 0 | 0 |
| **IC50 <50ug/ml** | 13 | 47 | 50 |
| **IC50 <10ug/ml** | 7 | 39 | 47 |
| **IC50 <1.0ug/ml** | 1 | 6 | 35 |
| **IC50 <0.1ug/ml** | 0 | 0 | 12 |
| **IC50 <0.01ug/ml** | 0 | 0 | 0 |
| **Median IC50** | 9.3100 | 3.1400 | 0.3030 |
| **Geometric Mean** | 6.3105 | 3.2488 | 0.8981 |

| | G70W-2E9 | J601-1B2 | VRC34.01 |
|---|---|---|---|
| **# Viruses** | 208 | 208 | 208 |
| **IC80 <50ug/ml** | 7 | 81 | 76 |
| **IC80 <10ug/ml** | 2 | 39 | 67 |
| **IC80 <1.0ug/ml** | 1 | 3 | 48 |
| **IC80 <0.1ug/ml** | 0 | 0 | 1 |
| **IC80 <0.01ug/ml** | 0 | 0 | 0 |
| **IC80 <50ug/ml** | 3 | 39 | 37 |
| **IC80 <10ug/ml** | 1 | 19 | 32 |
| **IC80 <1.0ug/ml** | 0 | 1 | 23 |
| **IC80 <0.1ug/ml** | 0 | 0 | 0 |
| **IC80 <0.01ug/ml** | 0 | 0 | 0 |
| **Median IC80** | 24.1000 | 10.1000 | 0.8415 |
| **Geometric Mean** | 12.0444 | 9.1101 | 1.6879 |

**C**

| mAb ID. | probe | Binding in ELISA | VH_call | VH_pct_SHM | JH_call | Isotype | CDRH3_AA | CDRH3 net charge | CDRH3_length | LightChain Type | VKL_call | VKL_pct_SHM | JKL_call | CDRL3_AA |
|---|---|---|---|---|---|---|---|---|---|---|---|---|---|---|
| J601-1B2 | BG505+JG2+ | BG505+459C+FP9+T250+ | IGHV4-AGR*01 | 19.5 | IGHJ5-1*01 | IGHG4*03_F_CH1 | CSSTYSWNNVWFDQW | -1 | 13 | Kappa | IGKV1-ABZ*01 | 12.2 | IGKJ4*01 | CQQRYSFPITF |
| G70W-2E9 | BG505+JG2+ | BG505+459C+FP9+T250+ | IGHV4-AGU*01 | 15.9 | IGHJ5-1*01 | IGHG4*03_F_CH1 | CLTRYNFYFGYRPNNRIDVW | 2 | 18 | Kappa | IGKV1-ACY*01 | 7.7 | IGKJ4*01 | CLQDDSFPLTF |

**D**

FP-specific antibody sequences.

| mAb ID. | V-GENE and allele | V-REGION identity % (nt) | J-GENE and allele | CDR-IMGT lengths | AA JUNCTION | V-GENE and allele | V-REGION identity % (nt) | J-GENE and allele | CDR-IMGT lengths | AA JUNCTION |
|---|---|---|---|---|---|---|---|---|---|---|
| G70W-2E9 | IGHV4S13*01 F | 89.80% | IGHJ5-1*01 F | [10.8.18] | CLTRYNFYFGYRPNNRIDVW | IGKV1-94*01 F | 93.55% | IGKJ4*01 F | [6.3.9] | CLQDDSFPLTF |
| J601-A6 | IGHV4S13*01 F | 84.35% | I IGHJ4*01 F | [10.8.13] | CSSTYSWNNVWFDQW | IGKV1-38*02 F | 90.32% | IGKJ4*01 F | [6.3.9] | CQQRYSFPITF |
| J601-1B2 | IGHV4S13*01 F | 83.67% | IGHJ4*01 F | [10.8.13] | CSSTYSWNNVWFDQW | IGKV1-38*02 F | 89.61% | IGKJ4*01 F | [6.3.9] | CQQRYSFPITF |
| J618-P1G4 | IGHV3-16*01 F | 90.72% | IGHJ4*01 F | [8.9.15] | CTTRHYNLGTGHFFDYW | IGKV2-60*01 F | 93.88% | IGKJ2*01 F | [11.3.9] | CMQGLQIPHTY |
| J618-P2E7 | IGHV3-16*01 F | 93.88% | IGHJ5-2*01 F | [8.10.15] | CTTRHYNLWTGHFFDYW | IGKV2-60*01 F | 93.20% | IGKJ2*01 F | [11.3.9] | CMQGLHTPPSF |
| J618-P2H6 | IGHV4-65*01 F | 91.41% | IGHJ4*01 F | [9.8.10] | CARESPLDFDYW | IGKV2-64*01 F | 94.56% | IGKJ1*01 F | [11.3.9] | CGQGIVLPRTF |
| J618-P2A12 | IGHV5-20*01 F | 90.97% | IGHJ5-1*02 F | [8.8.22] | CAIVRAGSYRDQYGHYYNSWFDVW | IGKV1-25*01 F | 86.38% | IGKJ2*01 F | [6.3.9] | CQHGYSNPYTF |
| K001-B9 | IGHV3-54*02 F | 93.75% | IGHJ4*01 F | [8.8.15] | CARDQSYYFDSGFPDFW | IGKV3-42*02 F | 94.27% | IGKJ3*01 F | [6.3.9] | CQKYDNSPFSF |
| K001-B8 | IGHV4-106*01 F | 91.41% | IGHJ4*01 F | [9.8.17] | CSGRRLVYYNLWNSPTDFW | IGKV1-16*01 F | 94.27% | IGKJ1*01 F | [6.3.9] | CQQYNSPPWTF |
| K001-A10 | IGHV4-93*01 F | 90.03% | IGHJ4*01 F | [9.8.11] | CARVVDGSHFEYW | IGKV1-21*01 F | 97.49% | IGKJ2*01 F | [6.3.10] | CQQYNSAPLYSF |

**Figure S4. FP- specific antibodies sorted from week 68 PBMC, related to Figure 4. (A)** Screening of FP-specific mAbs with three virus panel. mAbs isolated from G1, G2 and G3 NHPs are in black, blue and green fonts, respectively. (**B**) Neutralization of two FP-specific mAbs on 208 virus panel, as shown the number of viruses that were inhibited by 50% (IC50) or 80% (IC80) by the tested mAbs, and median IC50 and IC80 titers for the neutralized viruses. (**C**). Probes, sequence and binding profiles of two FP-specific mAbs that were tested in 208 virus panel. (**D**) Antibody sequence alignment of FP-specific mabs isolated from this study.

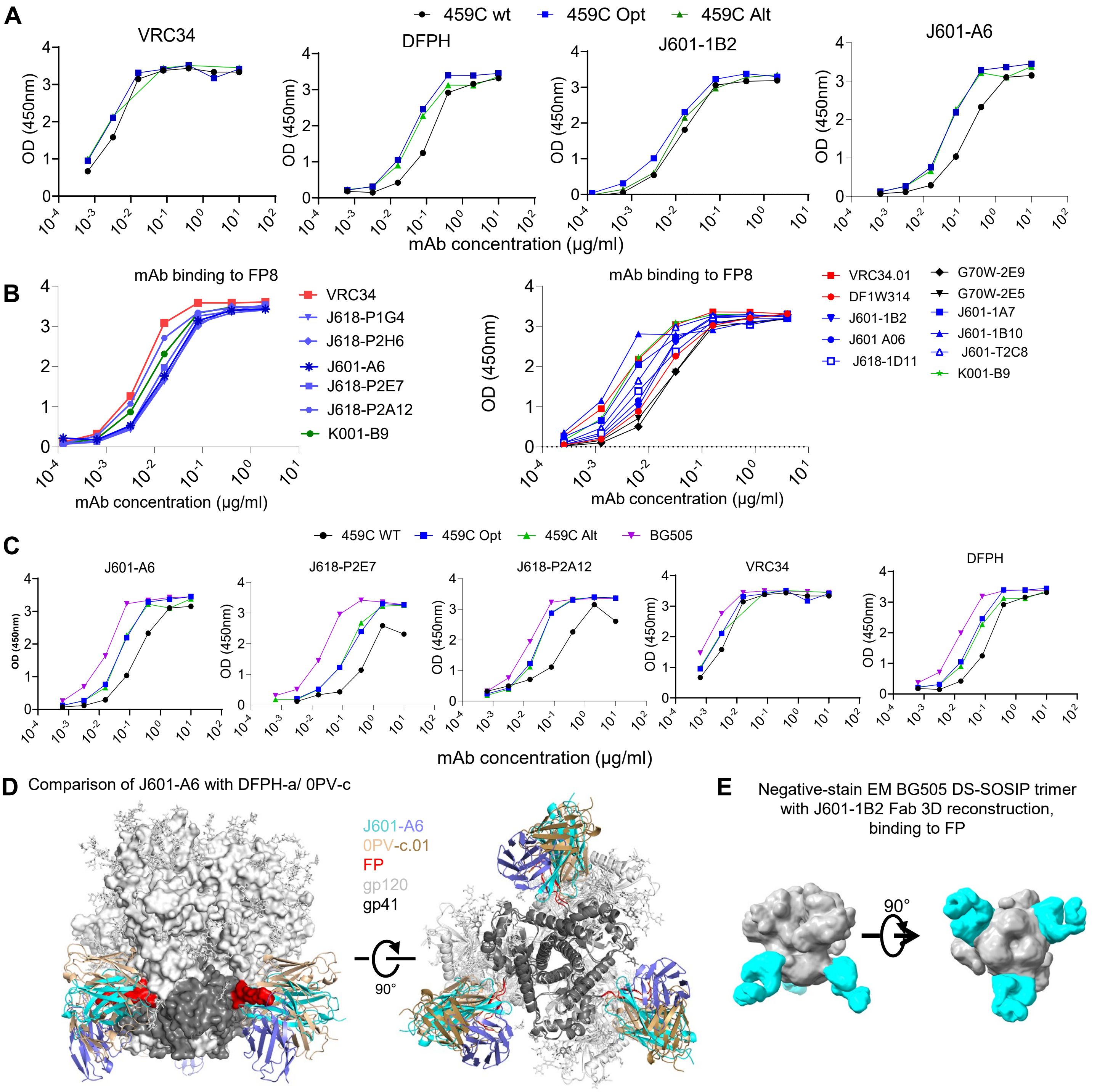

**Figure S5. Antigenicity and structural analysis of FP-specific mAbs isolated from the immunized NHPs reveals a binding mode similar to that of the previously identified DFPH-a class antibodies, related to Figure 4.** (**A, B**) mAbs binding to 459C WT, OPT and ALT trimers (**A**) and FP8 peptide (**B**); (**C**) Comparison of FP-specific mAbs binding to autologous and heterologous trimers. (**D**) Comparison of the binding mode of J601-A6 with that of 0PV-c.01 (PDB: 6NF2), an antibody of the DFPH-a class. The two structures were aligned by one of the gp120 subunits. On the left, Env trimer is shown a surface representation; on the right, both Env trimer and antibodies are shown in cartoon representation, with glycans shown in sticks. (**E**) Negative-stain EM of BG505 DS-SOSIP trimer with J601-1B2 Fab 3D reconstruction, binding to FP site of the trimer.

**A**

| Sort method | mAb specificity | Probes | samples | Plasmid synthesis | No. of synthesized mAbs |
|---|---|---|---|---|---|
| 1st round of sort | FP-specific | 459c V2 MIX APC (WT, OPT, ALT, 1:1:1) +FP9-Peg12-PE, double positives | wk68 | 1st and 2nd batch mAbs | 53+27 |
| | V2-specific | T250⁺ +T250.K169E⁺ | wk68 | 1st and 2nd batch mAbs | 53+27 |
| 2nd round of sort | V2-specific | Cap256+K169E- | wk68 | 3rd and 4th batch mAbs | 27+10 |
| | FP-specific | BG505-APC+JG2-PE+ | wk68 | 3rd and 4th batch mAbs | 27+10 |
| 3rd round of sort | V2-specific | T250.wt-APC+ T250.AF700+ T250.K169E- | wk68, wk132, wk156 | 5th batch | 16 |
| 4th round of sort | V2 or FP -specific | H703, CT184, T250 trimer probes, any 2 trimer+ regardless of K169E profile | wk68, wk132, wk156 | 6th batch | 26 |
| 5th round of sort | V2 or FP | CT184, T250, T250.V1V2.gp120, 459C OPT, 459C WT, any 2-probe positive | wk132, wk68 | 7th batch | 23 |
| 6th round of sort | V2 or FP | H703, CT184, ZM233, any 2-probe positive | Wk132, wk68 | | |

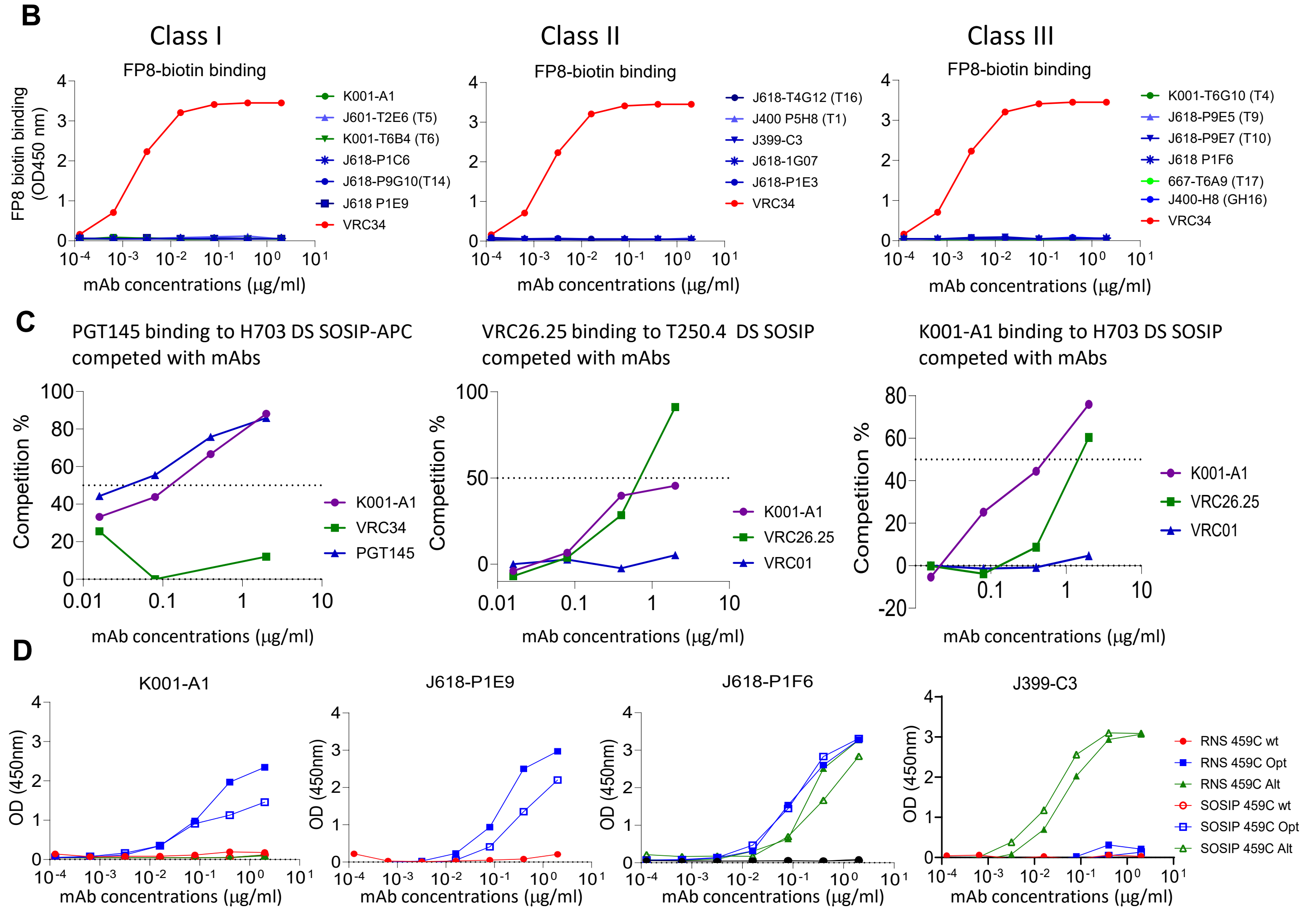

**Figure S6. V2-apex and FP-specific antibodies were sorted from immunized NHPs, related to Figure 4-6. (A).** Summary of sort method, probes and mabs synthesized from PBMCs at wk68, wk132 or wk156 after immunization. **(B).** V2 apex-specific mAbs do not bind to FP peptide detected with ELISA. (**C**) V2-specific mAbs isolated from this study compete with human V2 apex bNabs on env trimer binding. (**D)** Binding of V2-specific mabs to 459C WT, OPT and ALT DS SOSIP trimers and RnS stabilized trimers in ELISA.

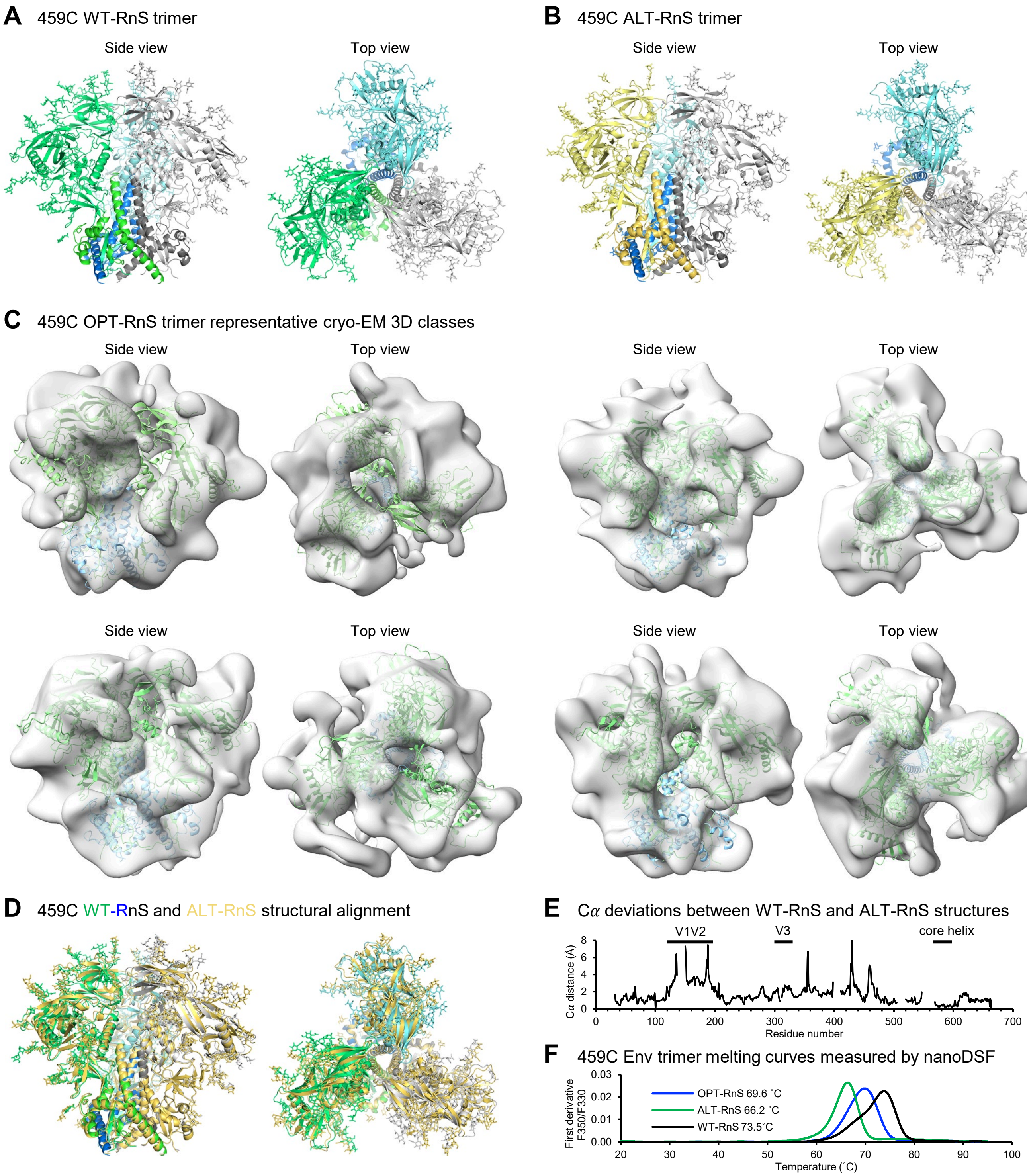

**Dats S1.1. Cryo-EM structural analysis of RnS stabilized 459C WT, OPT, and ALT Env trimers, related to Figure 1**

(A) Cryo-EM structure of 459C WT-RnS trimer shown as cartoon representation. The three protomers are colored individually in green, blue, and gray. (B) Cryo-EM structure of 459C ALT-RnS trimer shown as cartoon representation. The three protomers are colored individually in yellow, blue, and gray. (C) Cryo-EM 3D reconstruction maps of 459C OPT-RnS trimer particles, superimposed with the cryo-EM structure of 459C WT-RnS. (D) Structural superposition of 459C WT-RnS and ALT-RnS structures, based on all alignable Cα atoms. WT-RnS is colored green, blue and gray; ALT-RnS is colored yellow. The structures were aligned in Pymol. The apex region shows some shifts between the structures, with the ALT structure expanding outwards. The gp41 core helices superimpose especially well. (E) Deviation plot between Cα atoms of the aligned 459C WT-RnS and ALT-RnS structures. (F) Protein melting curves measured by nanoDSF. The protein concentrations were 0.9 mg/ml in 20 mM Na citrate pH 6.0, 75 mM NaCl and 5% sucrose. The samples were heated at 1 ˚C/min from 20 to 95 ˚C.

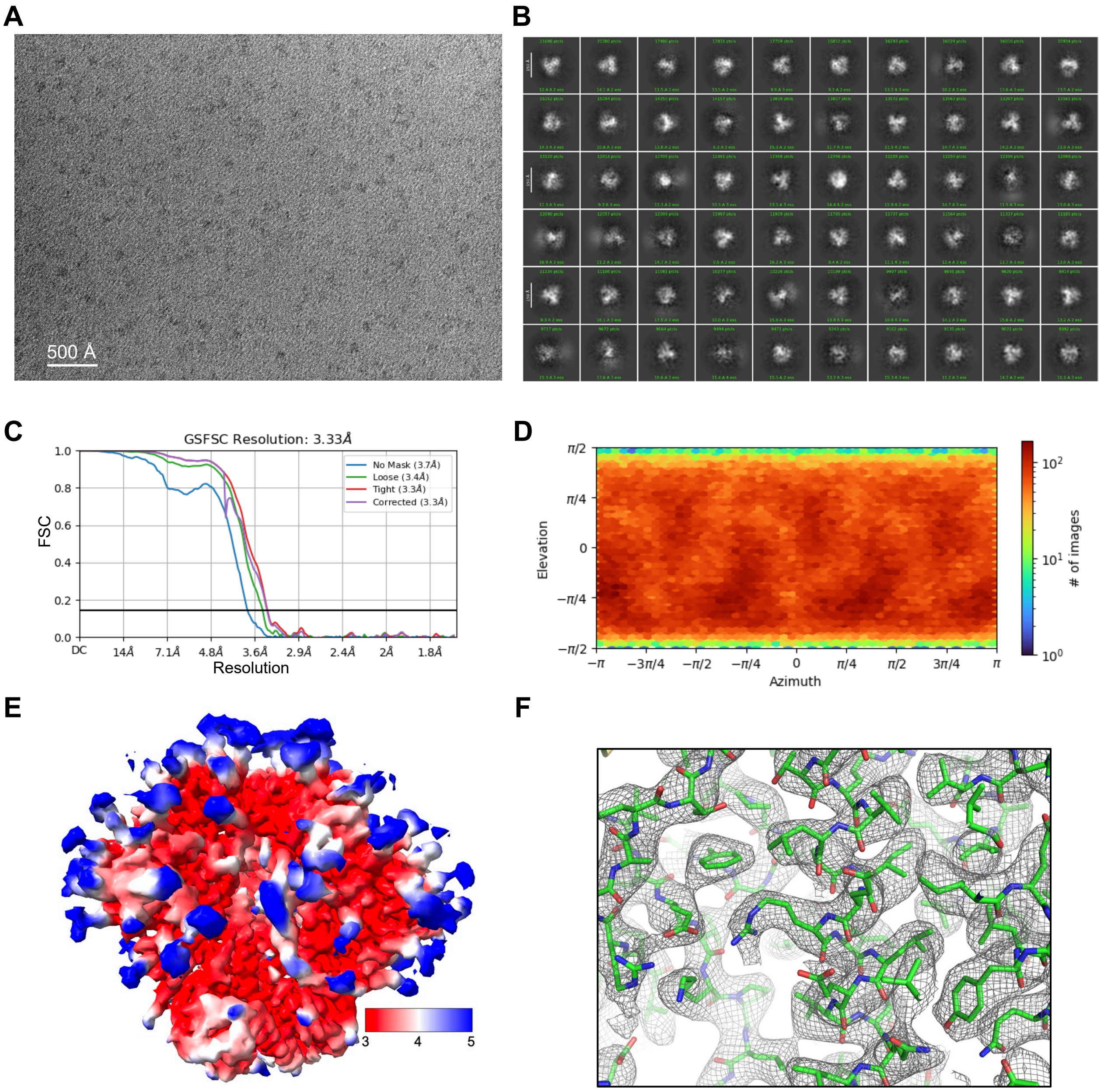

**Data S1.2. Cryo-EM validation for 459C-WT-RnS, related to Figure 1**

(A) Representative micrograph. (B) 2D classes. (C) Fourier shell correlation (FSC) curves showing gold-standard FSC resolution at 3.33 Å. (D) Orientation of all particles used in the final refinement shown as a heatmap. (E) Local resolution as indicated by colors on the final 3D reconstruction density map. (F) A section of representative density of cryoSPARC sharpened map contoured at 6$\sigma$.

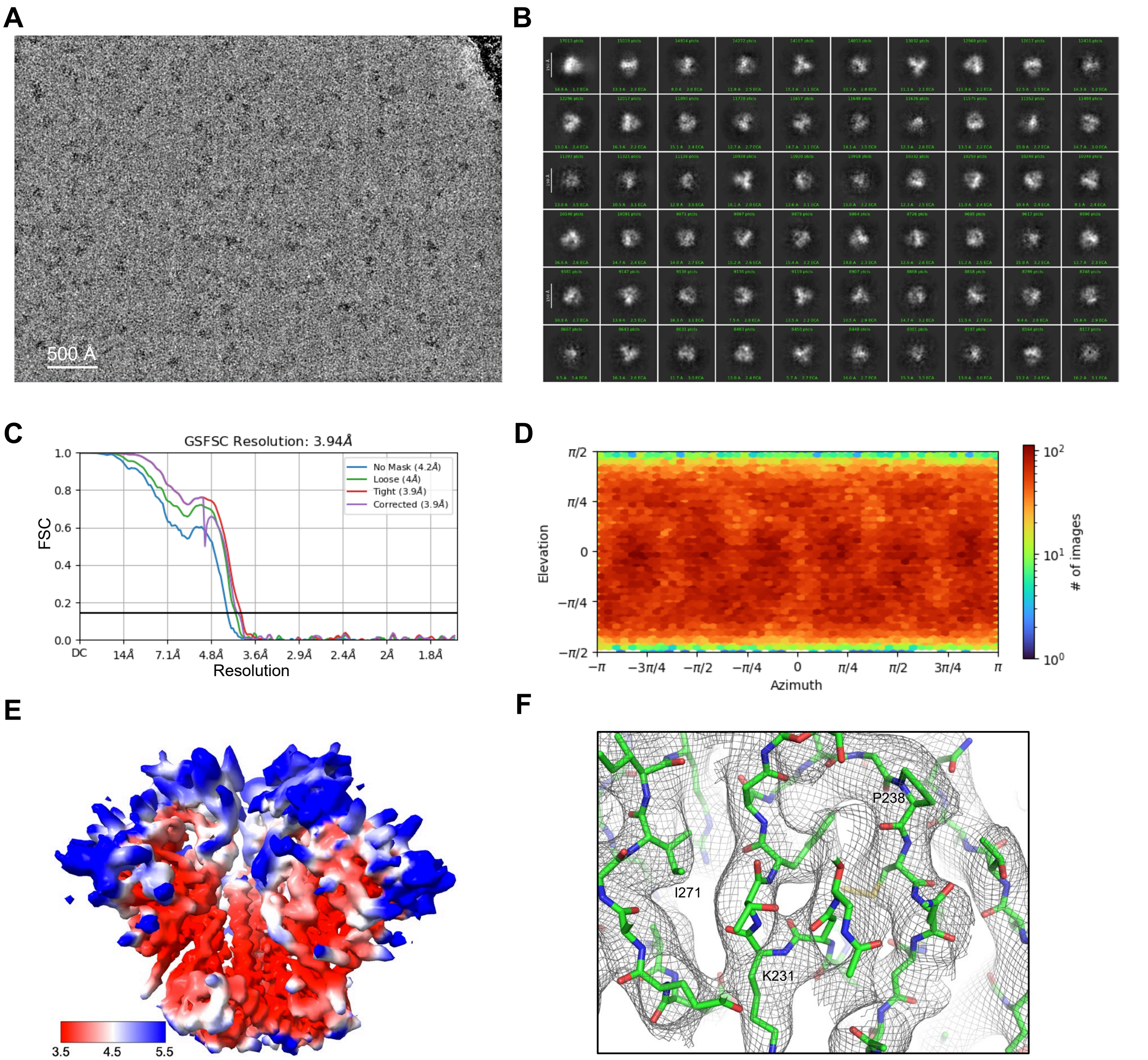

**Data S1.3. Cryo-EM validation for 459C-ALT-RnS, related to Figure 1**
(A) Representative micrograph. (B) 2D classes. (C) Fourier shell correlation (FSC) curves showing gold-standard FSC resolution at 3.94 Å. (D) Orientation of all particles used in the final refinement shown as a heatmap. (E) 3D reconstruction density map with local resolution indicated by colors. . (F) A section of representative density of cryoSPARC sharpened map contoured at 6$\sigma$.

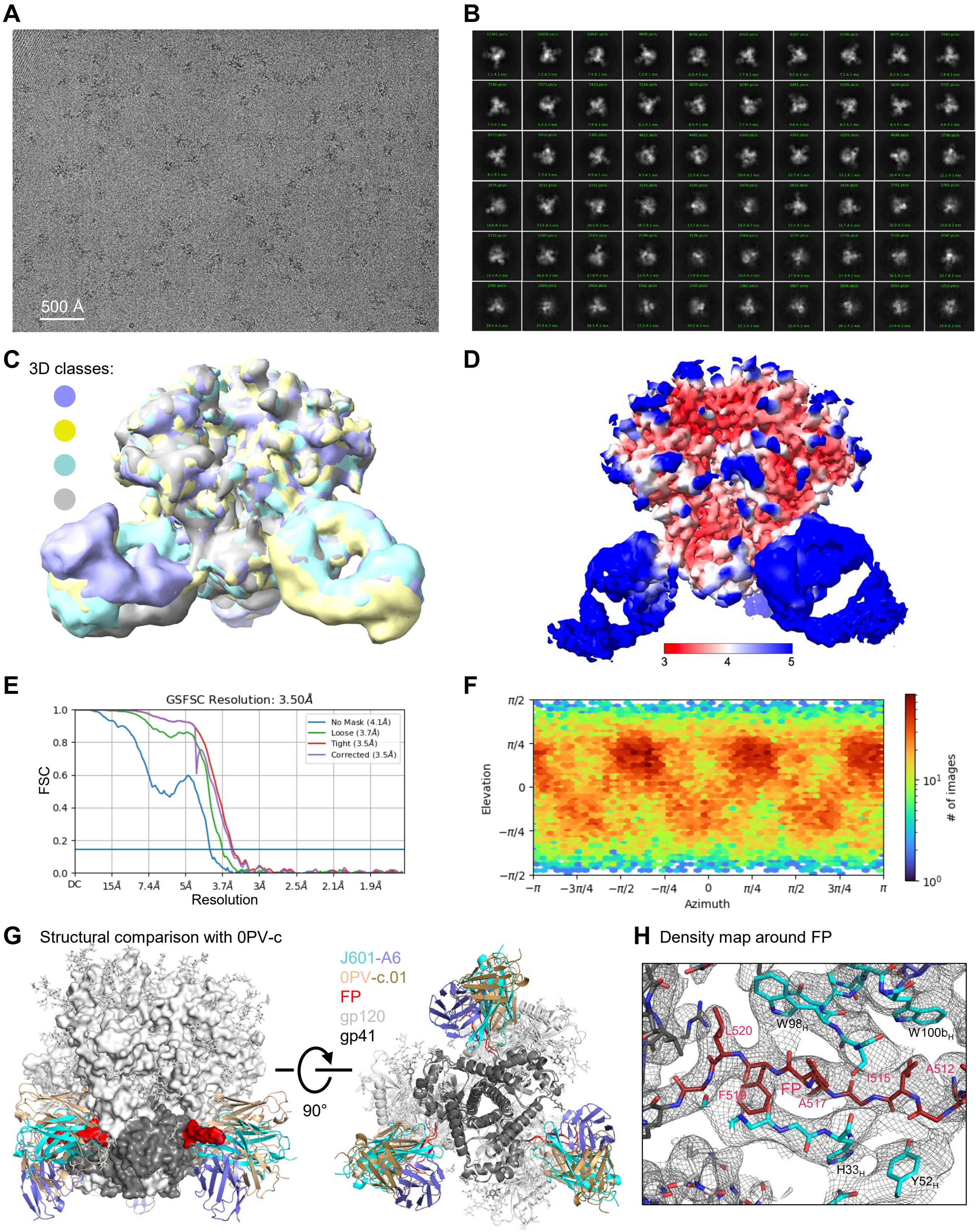

**Data S1.4. Cryo-EM validation for J601-A6 in complex with BG505 DS-SOSIP, related to Figure 4**

(A) Representative micrograph. (B) 2D classes. (C) Multiple 3D classes with varying antibody orientations. (D) Local resolution of one of the 3D classes with C3 symmetry at a nominal resolution of 3.50 Å. (E) Fourier shell correlation (FSC) curves showing gold-standard FSC resolution at 3.50 Å. (F) Orientation of particles used in the final refinement shown as a heatmap. (G) Comparison of the binding mode of J601-A6 with that of 0PV-c.01 (PDB: 6NF2), an antibody of the DFPH-a class. The two structures were aligned by one of the gp120 subunits. Left, Env trimer is shown as a surface representation; right, both Env trimer and antibodies are shown in cartoon representation, with glycans shown as sticks. (H) A section of density map around the N-terminal segment of FP interacting with J601-A6 heavy chain, contoured at 5.5$\sigma$.

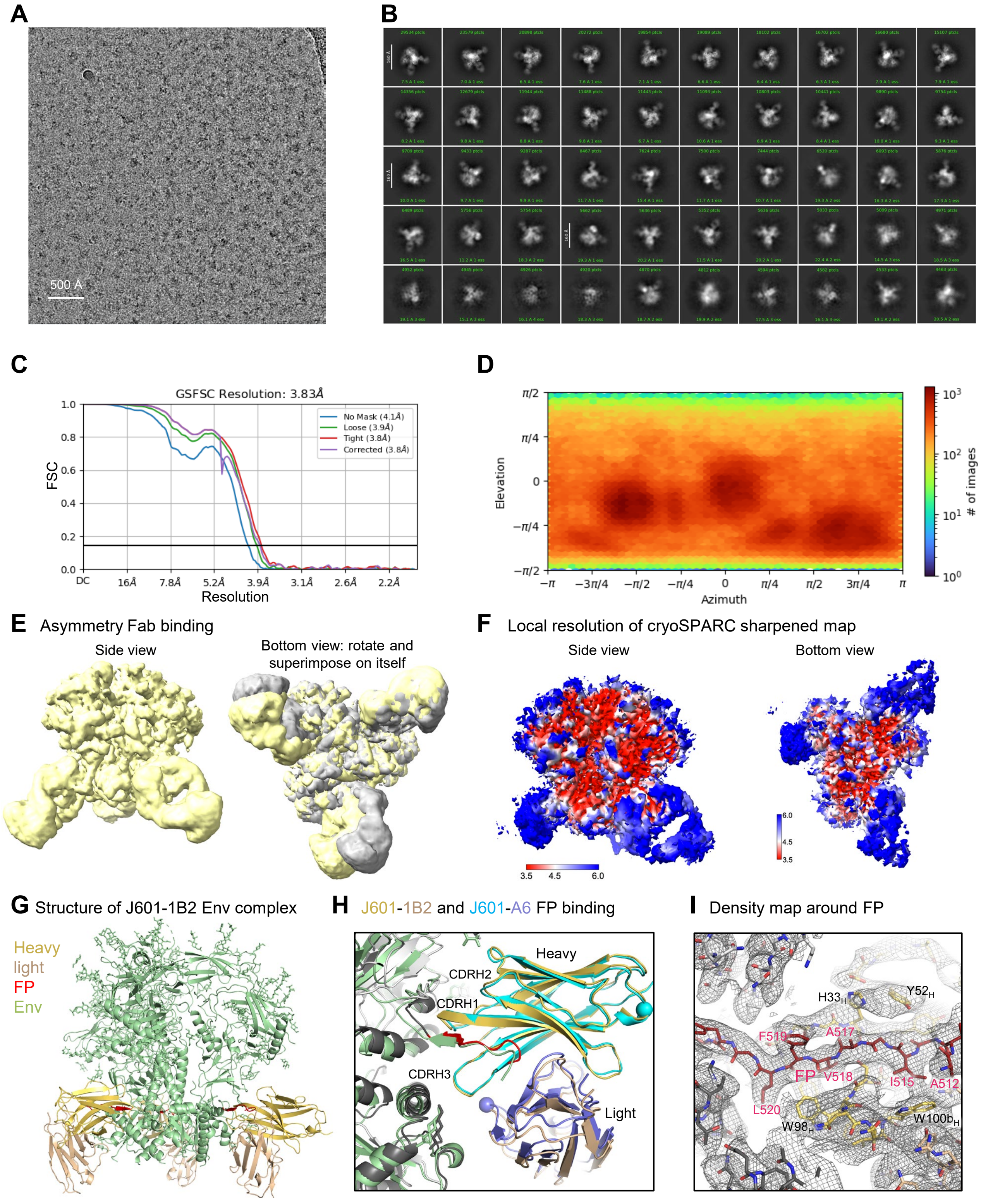

**Data S1.5. Cryo-EM validation for J601-1B2 in complex with BG505 DS-SOSIP, related to Figure 4**

(A) Representative micrograph. (B) 2D classes. (C) Fourier shell correlation (FSC) curves showing gold-standard FSC resolution at 3.83 Å refined with C1 symmetry. (D) Orientation of particles used in the final refinement shown as a heatmap. (E) 3D reconstruction map showing asymmetric binding of J601-1B2 Fab at the FP site. (F) Local resolution of cryoSPARC sharpened map. (G) Overall structure of refined model of J601-1B2 binding to BG505 DS-SOSIP. (H) Comparison of the binding mode of J601-1B2 with that J601-A6 by aligning the heavy chains, revealing identical binding interactions with FP. Spheres indicate the four different residues between them, two of them behind the strands between CDRs H1 and H2. (I) A section of density map around the N-terminal segment of FP interacting with J601-1B2 heavy chain, contoured at 6$\sigma$.

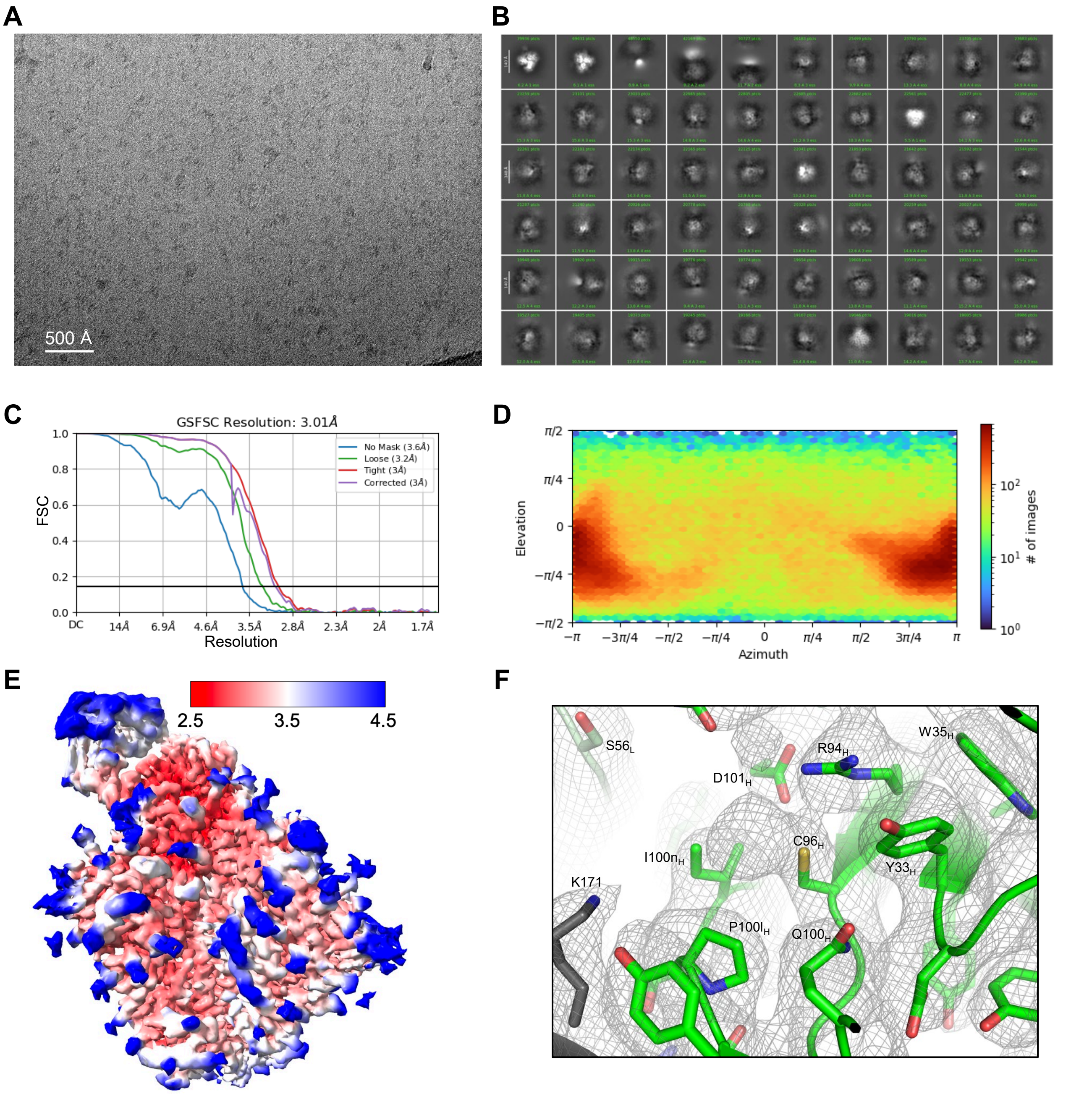

**Data S1.6. Cryo-EM validation for K001-A1 in complex with 459C-OPT, related to Figure 7**
(A) Representative micrograph. (B) 2D classes. (C) Fourier shell correlation (FSC) curves showing gold-standard FSC resolution at 3.01 Å. (D) Orientation of all particles used in the final refinement shown as a heatmap. (E) 3D reconstruction density map at 3.01 Å from non-uniform refinement with C1 symmetry with local resolution indicated by colors, showing antibody Fab binding at Env apex. (F) Representative density map. A section of density map around C96 of the K001-A1 heavy chain is shown at a contour level of 6 $\sigma$. The single unpaired cysteine C96 has extra density extending beyond the S$\gamma$ atom, indicating it is likely modified. It is unclear what modification at this resolution and is therefore not modeled.

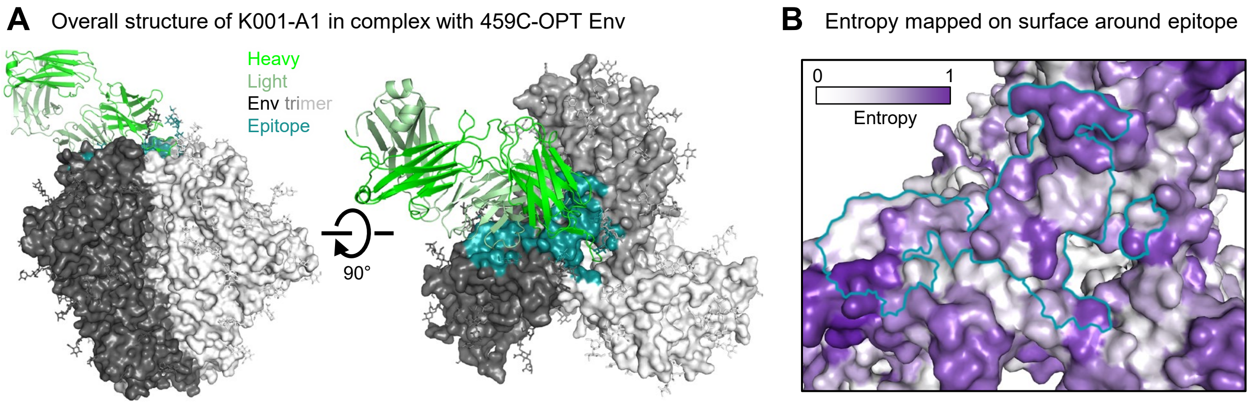

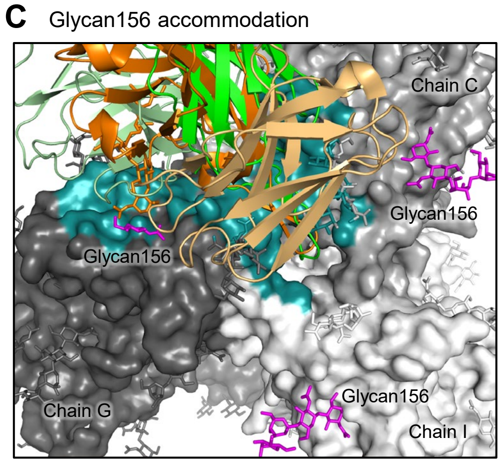

**D** PG9 and VRC26.25 binding Env protein involves only heavy chain

| HIV-1 Env ApexGT3.N130 in complex with PG9 – 7t77 | | | | HIV-1 Env CAP256.wk34.c80 SOSIP.RnS2 in complex with CAP256-VRC26.25 – 6vtt | | | |
|---|---|---|---|---|---|---|---|
| Antibody heavy chain | | | | Antibody heavy chain | | | |
| | Epitope region | BSA(Å²) | Fab region | | Epitope region | BSA(Å²) | Fab region |
| Chain C | C-strand 169 | 56.5 | CDRH3 | Chain F | C-strand 169 | 35.7 | CDRH3 |
| | V1V2 184-186 | 131.9 | CDRH1 | | Apex hole 162,166,167 | 109.2 | CDRH3 |
| | | | | | 123,124,128,190, | 60.2 | CDRH3 |
| | | | | | 312,313,315 | 28.2 | CDRH3 |
| | | | | | Glycan 160 | 41.0 | CDRH3 |
| Chain E | Apex hole 165-167 | 71.0 | CDRH3 | Chain G | C-strand 168-171 | 307.0 | CDRH3 |
| | V1V2 127,158,160 | 46.8 | CDRH3 | | Apex hole 160-167 | 228.5 | CDRH3 |
| | C-strand 168-173 | 447.7 | CDRH3 | | 123,124,127 | 53.6 | CDRH3 |
| | Glycan 160 | 402.9 | CDRH3 | | 313,315 | 7.4 | CDRH3 |
| | Glycan 156 | 247.7 | CDRH2,3 | | Glycan 130 | 108.4 | CDRH1,3 |
| | | | FRH3 | | Glycan 156 | 29.1 | CDRH3 |
| | | | | | Glycan 160 | 161.8 | CDRH3 |
| Chain A | C-strand 169 | 49.7 | CDRH3 | Chain E | C-strand 169 | 39.4 | CDRH3 |
| | Apex hole 166-167 | 45.8 | CDRH3 | | Apex hole 123 | 24.9 | CDRH3 |
| | | | | | 162,166,167 | 161.9 | CDRH3 |
| Total | | 1500.0 | | | | 1396.3 | |
| Chain E | Glycan160 – Fab L | 205.8 | CDRL1,2,3 | Chain G | Glycan156 – Fab L | 29.3 | CDRL2 |

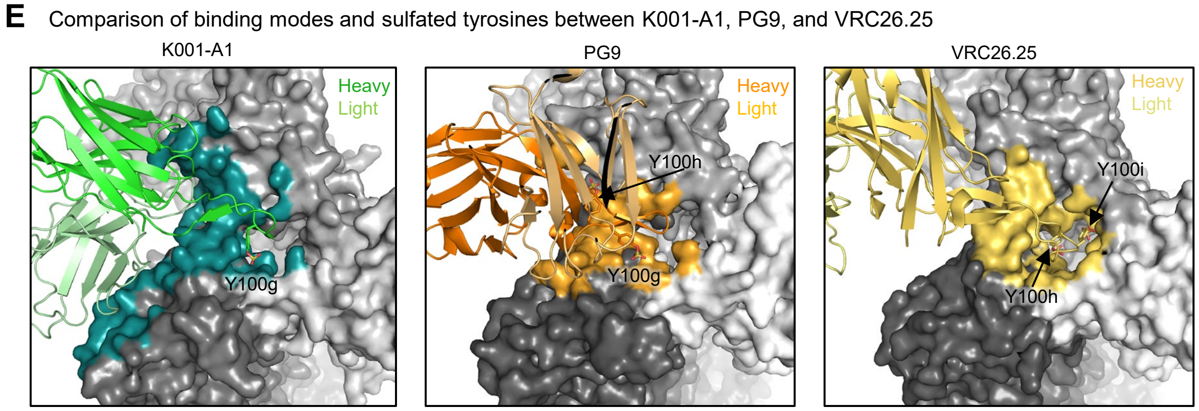

**Data S1.7. K001-A1 has a binding mode similar to that of PG9, but with additional light chain interactions, related to Figure 7.**

(A) Cryo-EM structure of K001-A1 Fab in complex with 459C-OPT-RnS Env trimer at 3.0 Å. Fab heavy chain is colored bright green and light chain in pale green; Env trimer is shown in surface representation, with different shade of gray for each protomer and color teal for the epitope. (B) Shannon entropy mapped on the 459C-OPT Env surface around the K001-A1 epitope. The entropy values of each residue is colored white to purple, with the normalized entropy value of 1 in dark purple. The entropy values were calculated with a curated alignment of year 2022 HIV-1 M group Env of 6,446 sequences (https://www.hiv.lanl.gov/content/sequence/NEWALIGN/align.html). (C) Superposition of PG9 complex with K001-A1 complex for glycan156 binding interactions. 459C-Opt Env trimer is shown as surface representation with different shades of gray for the three protomers and magenta for glycan156. K001-A1 Fab is bright green for heavy chain and pale green for light chain. PG9 Fab is orange for heavy chain and light orange for light chain. Glycan 156 on the dark gray protomer (left) is mostly disordered. It would clash with K001-A1 light chain if it adopted the same conformation as that in the PG9 complex, which interacts favorably with PG9 heavy chain. The other two glycan156 (top right and bottom) adopt the same conformation as those in the PG9 complex. (D) PG9 and VRC26.25 binding analysis by PISA. Light chains are involved only in minimal binding to glycans. (E) Comparison of overall binding modes between K001-A1, PG9, and VRC26.25 in complexes with Env trimers. All of them involved sulfated tyrosines. K001-A1 has one sulfated tyrosine, which is similar in binding location to one of the two sulfated tyrosines in PG9. Both sulfated tyrosines in VRC26.25 bind deeply in the apex hole.

**Data S1.8. Table of cryo-EM data and refinement statistics, related to Figures 1, 4, and 7**

| | 459C-WT-RnS | 459C-ALT-RnS | J601-A6 BG505 DS-SOSIP | J601-1B2 BG505 DS-SOSIP | K001-A1 459C-OPT |
|---|---|---|---|---|---|
| **EMDB ID** | EMD-71781 | EMD-71782 | EMD-71767 | EMD-71766 | EMD-71772 |
| **PDB ID** | 9PQ2 | 9PQ3 | 9PNN | 9PNI | 9PNU |
| Data collection | | | | | |
| Microscope | FEI Titan Krios | FEI Titan Krios | FEI Titan Krios | FEI Titan Krios | FEI Titan Krios |
| Voltage (kV) | 300 | 300 | 300 | 300 | 300 |
| Electron dose ($e^-/Å^2$) | 58 | 58 | 58 | 40 | 58 |
| Detector | Gatan K3 BioQuantum | Gatan K3 BioQuantum | Gatan K3 BioQuantum | Direct Electron Apollo | Gatan K3 BioQuantum |
| Pixel size (Å) | 0.825 | 0.825 | 0.83 | 0.505/1.01 | 0.825 |
| Defocus range (µm) | -0.8/-2.0 | -0.8/-2.0 | -0.8/-2.0 | -1.0/-2.5 | -0.8/-2.0 |
| Reconstruction | | | | | |
| Software | cryoSPARC v4.4.1 | cryoSPARC v4.4.1 | cryoSPARC v4.2.1 | cryoSPARC v4.4.1 | cryoSPARC v4.4.1 |
| Particles | 207,797 | 158,204 | 46,939 | 806,645 | 218,268 |
| Symmetry | C3 | C3 | C3 | C1 | C1 |
| Resolution (Å) ($FSC_{0.143}$) | 3.33 | 3.94 | 3.50 | 3.83 | 3.01 |
| Refinement | | | | | |
| Software | Phenix 1.21 | Phenix 1.21 | Phenix 1.21 | Phenix 1.21 | Phenix 1.21 |
| Protein residues | 1758 | 1728 | 2442 | 2442 | 2197 |
| Ligands | BMA: 36; NAG: 111; MAN: 24 | BMA: 30; NAG: 105; MAN: 18 | BMA: 36; NAG: 105; MAN: 21 | BMA: 32; NAG: 103; MAN: 22 | BMA: 31; NAG: 110; MAN: 20 |
| CC (box) | 0.82 | 0.85 | 0.82 | 0.86 | 0.81 |
| CC (mask) | 0.77 | 0.82 | 0.81 | 0.82 | 0.87 |
| R.m.s. deviations | | | | | |
| Bond lengths (Å) | 0.004 | 0.004 | 0.002 | 0.004 | 0.004 |
| Bond angles (°) | 0.906 | 0.858 | 0.534 | 0.759 | 0.632 |
| Validation | | | | | |
| Molprobity score | 1.51 | 1.25 | 1.51 | 1.34 | 1.50 |
| Clash score | 3.00 | 3.50 | 4.37 | 1.57 | 4.53 |
| Rotamer outliers (%) | 1.15 | 0.78 | 0.42 | 0.61 | 1.02 |
| Ramachandran | | | | | |
| Favored regions (%) | 94.46 | 97.53 | 95.75 | 93.12 | 96.11 |
| Allowed regions (%) | 5.54 | 2.47 | 4.25 | 6.88 | 3.89 |
| Disallowed regions (%) | 0 | 0 | 0 | 0 | 0 |

**Data S1.9 Buried surface area of epitope residues in the K001-A1 459C-Opt complex, related to Figure 7**
Normalized entropy values were calculated with Shannon Entropy-One (https://www.hiv.lanl.gov/content/sequence/ENTROPY/entropy_one.html) on the curated alignment of year 2022 HIV-1 M group Env of 6,446 sequences (https://www.hiv.lanl.gov/content/sequence/NEWALIGN/align.html). A residues is highlighted in yellow when different among all three constructs or the one different from the other two. Buried surface area (BSA) is calculated with PISA. Chains G, C, and I are the three gp120 subunits; chains H and L and the heavy and light chains of Fab.

| Residue | | Amino acid in | | | BSA($Å^2$) between chains | | | | |
|---|---|---|---|---|---|---|---|---|---|
| number | Entropy | Opt | Alt | Wt | G-H | G-L | C-H | C-L | I-H |
| Residues in and around the apex hole | | | | | | | | | |
| 124 | 0.0125 | P | P | P | 4.8 | | | | |
| 127 | 0.0327 | V | V | V | 15.55 | | 4.52 | | |
| 128 | 0.107 | T | T | T | | | 30.12 | | |
| 130 | 0.430 | E | D | N | | | 1.6 | | |
| 160 | 0.153 | N | N | N | 9.72 | | 17.32 | | |
| 161 | 0.540 | M | I | A | 1.83 | | | | |
| 162 | 0.140 | T | T | T | 37.09 | | | | |
| 163 | 0.176 | T | T | T | 0.83 | 0.5 | | | |
| 164 | 0.701 | E | S | E | | 37.92 | | | |
| 165 | 0.496 | L | V | I | 4.44 | 2.68 | | | |
| 166 | 0.455 | R | K | R | 3.88 | | | | 52.88 |
| 167 | 0.290 | D | G | D | 79.76 | | | | 17.43 |
| 168 | 0.205 | K | K | R | 82.09 | 20.41 | | | |
| 169 | 0.754 | K | R | K | 101.4 | | 28.88 | | 24.86 |
| 170 | 0.542 | K | Q | K | 57.53 | 61.86 | | | |
| 171 | 0.458 | K | Q | E | 77.74 | 7.59 | | | |
| 172 | 0.550 | V | E | M | 0.41 | 10.61 | | | |
| 173 | 0.393 | S | H | Y | | 18.99 | | | |
| 175 | 0.222 | L | L | L | | 2.66 | | | |
| 182 | 0.295 | V | V | V | | | | 3.52 | |
| 183 | 0.408 | P | P | P | | | | 11.91 | |
| 184 | 0.364 | L | L | L | | | 10.92 | 47.79 | |
| 185 | 0.606 | N | N | D | | | 24.97 | 3.68 | |
| 186 | 0.667 | K | K | K | | | 37.14 | 78.96 | |
| 187 | 0.670 | N | N | S | | | 96.92 | | |
| 189 | 0.649 | R | R | N | | | 19.5 | | |
| 190 | 0.669 | Q | Q | R | | | 31.86 | | |
| 192 | 0.341 | R | R | R | | | | 21.19 | |
| 197 | 0.0408 | N | N | N | | | | 12.6 | |
| 308 | 0.577 | R | R | R | | 47.44 | | | |
| Residues distal to the apex hole | | | | | | | | | |
| 133 | 0.504 | A | A | N | | 2.09 | | | |
| 134 | 0.819 | F | F | V | | 30.05 | | | |
| 135 | 0.814 | N | N | T | | 13.26 | | | |
| 136 | 0.900 | S | S | S | | 42.45 | | | |
| 303 | 0.0739 | T | T | T | | 26.09 | | | |
| 304 | 0.102 | R | R | R | | 0.48 | | | |
| 305 | 0.321 | K | K | K | | 31.9 | | | |
| 321 | 0.366 | N | N | N | | 50.53 | | | |
| 321A | 0.650 | D | E | D | | 56.07 | | | |
| 322 | 0.119 | I | I | I | | 15.45 | | | |
| 323 | 0.172 | I | I | I | | 42.01 | | | |
| 324 | 0.0213 | G | G | G | | 25.09 | | | |
| 325 | 0.231 | D | D | D | | 4.79 | | | |
| 326 | 0.0500 | I | I | I | | 6.8 | | | |
